# Dynamics and fermentation patterns of stool microbiota on simple carbon sources in *in vitro* batch cultures reveal dysbiosis in Crohn’s disease

**DOI:** 10.1101/2025.05.29.656803

**Authors:** Anna Detman-Ignatowska, Gabriele Schiro, Rafał Filip, Emilia Samborowska, Jakub Karczmarski, Kinga Jakubowska, Sara Jarmakiewicz-Czaja, Anna Williams, Nabahi Ramos Hickman, Daniel Laubitz, Anna Sikora

## Abstract

Crohn’s disease (CD)-associated dysbiosis is characterized by reduced microbial diversity, depletion of short-chain fatty acid (SCFA) producers, especially butyrate-forming taxa, and altered metabolic profiles. This study examined whether CD-related dysbiosis is reflected in the fermentation behavior of stool-derived microbial communities cultured in vitro on simple carbon substrates (glucose or acetate+lactate). Shotgun metagenomics and metabolomics revealed that CD-derived communities produced lower levels of butyrate (mainly *via* lactate and acetate), valerate, caproate, and propionate, and higher levels of ethanol and certain amino acids compared to healthy controls. These metabolic alterations aligned with compositional shifts, including a loss of beneficial commensals (e.g., *Coprococcus catus, Ruminococcus torques, Eubacterium rectale, Fusicatenibacter saccharivorans,* and *Faecalibacterium prausnitzii*) and an overrepresentation of CD-associated taxa, particularly *Escherichia coli.* Metabolic potential analysis revealed an enrichment of genes linked to ethanol and amino acid synthesis in CD-associated microbiotas, underscoring the metabolic adaptability of *E. coli.* Notably, acetate+lactate substrates supported the growth of healthy microbiota-associated bacteria, whereas glucose favored CD-associated taxa, highlighting disease-specific metabolic imbalances. These findings suggest that in vitro fermentation profiling may help distinguish CD-associated dysbiosis from a healthy microbiome and might support the development of microbiome-informed diagnostic or therapeutic approaches for CD.

**Graphical abstract:** 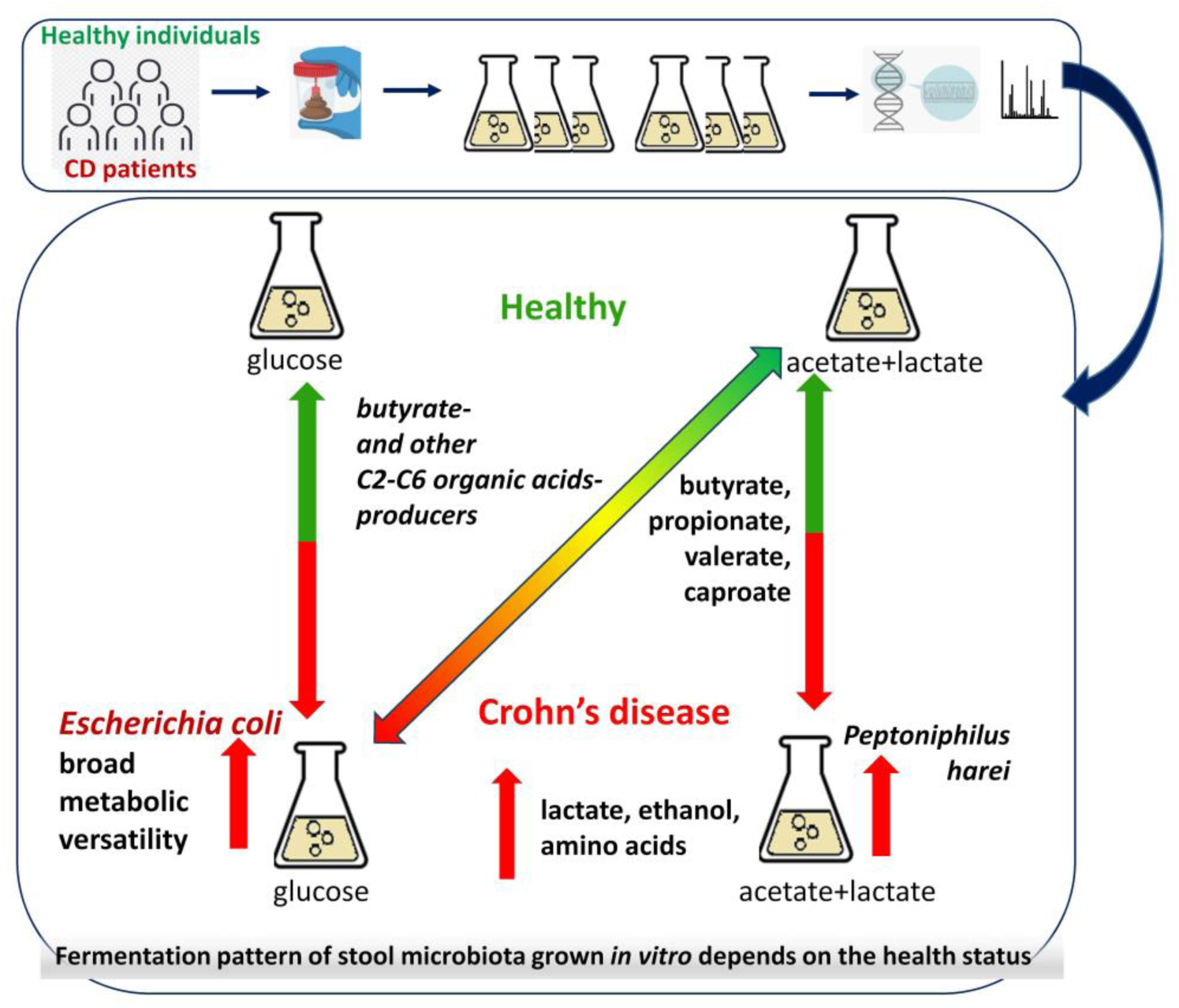

**A short abstract:** CD-related dysbiosis alters stool microbial fermentation, reducing butyrate, propionate, valerate, caproate, and increasing ethanol/amino acids *in vitro*. Fermentation tests may help distinguish CD-associated patterns, suggesting potential for future microbiome-based diagnostics.

## 1. Introduction

The crucial role of host-associated microbial communities in health and disease is becoming increasingly evident. The gut microbiome stands out among these communities due to its immense size (∼10¹³ microbial cells including bacteria, fungi, and archaea), extraordinary diversity, and functional complexity. It plays a central role in maintaining gastrointestinal (GI) tract homeostasis through processes such as the fermentation of dietary polysaccharides into C2-C6 organic acids like acetate, propionate, and butyrate, which serve as energy sources and regulators of host metabolism ^1–11^. The healthy human gut microbiome is dominated by bacteria: Firmicutes (recently reclassified as Bacillota) and Bacteroidetes (Bacteroidota), with smaller contributions from Proteobacteria (Pseudomonadota) and Actinobacteria (Actinomycetota) ^1–6^. However, disruptions in this intricate ecosystem, termed dysbiosis, have been implicated in a wide range of conditions, including inflammatory bowel disease (IBD), metabolic disorders, allergies, asthma, neurological diseases, and cancer ^5,12–18^. Considerable effort has been dedicated to identifying strategies to facilitate transitions from one microbial state to another to promote better health outcomes^18,19^. Inflammatory bowel disease, one of the most common and prevalent gut diseases, affecting up to 4% of the adult population, is mainly attributed to a Westernized lifestyle ^20^. It manifests primarily as ulcerative colitis (UC) or Leśniowski-Crohn’s disease, commonly known as Crohn’s disease (CD), the latter characterized by inflammation in the ileum, cecum, and colon. CD symptoms include abdominal pain and altered bowel habits, with untreated or late-diagnosed cases increasing the risk of colorectal cancer. Despite advances, the etiology of CD remains elusive, involving complex interactions between genetic, microbial, immunological, and environmental factors ^21,22^. A hallmark of CD is dysbiosis, characterized by reduced microbial richness and diversity, increased susceptibility to infections, and altered microbial metabolite profiles. These changes include an increased abundance of Proteobacteria (e.g., *Escherichia coli*) and Actinobacteria, a reduced Firmicutes-to-Bacteroidetes ratio, and a loss of butyrate producers, such as *Faecalibacterium prausnitzii* ^1,9,11–13,23–28^.

Dysbiosis-induced shifts in C2–C6 organic acids, essential for gut health, exacerbate inflammatory processes in CD. Decreased levels of acetate, butyrate and propionate are observed in stool samples of IBD patients, whereas lactate and pyruvate concentrations are increased ^1,7,9,29^. Metabolomic studies of feces, blood, urine and biopsied colonic samples from IBD patients have revealed changes in bile acid composition, lipid metabolism, amino acid metabolism and the synthesis of vitamins (especially of the B group) ^1,7,9,29–33^. Understanding the mechanisms underlying these shifts remains challenging, as invasive sampling methods hinder human studies and animal models fail to fully replicate human microbiome dynamics. Advances in *in vitro* cultivation offer an alternative, enabling the study of stool-derived microbial communities under controlled conditions. These systems have been used to investigate polysaccharide metabolism, probiotic effects, and dietary interventions ^34–36^ but rarely to assess the metabolic potential of stool microbiotas grown on simple carbon substrates, such as glucose, acetate, and lactate. Glucose, acetate, and lactate are relevant substrates in the gut environment. While glucose represents a primary product of dietary polysaccharide breakdown, acetate and lactate serve as intermediates in anaerobic fermentations and precursors for butyrate synthesis ^37,38^. Investigating the metabolic activity of stool microbiotas grown on these substrates could provide valuable insights into the microbial and metabolic signatures of health and disease. At the same time the experimental system does not require replicating gut conditions.

This study aimed to evaluate the dynamics and metabolic capacities of stool microbiotas from CD patients and healthy individuals grown in batch cultures with glucose or acetate+lactate as carbon sources. We hypothesized that the metabolic properties observed *in vitro* would reflect *in vivo* gut microbiome functionality, revealing genomic and ecological differences between healthy and CD-associated communities. By identifying taxa and metabolites indicative of these states, our findings might contribute to better diagnostic tools for CD, where the diagnosis can be delayed by years and silent cases remain undetected ^1,39^.

## 2. Materials and methods

### 2.1. Patients and justification of sample size

Stool samples were collected from patients with Crohn’s disease (CD) and healthy individuals at the Centre for Comprehensive Treatment of Inflammatory Bowel Diseases, Department of Gastroenterology, Clinical Regional Hospital No. 2 in Rzeszow, Poland. The CD group (n=7; CD1– CD7) included four males and three females aged 23–50 years. Clinical characteristics of the CD patients are summarized in Table S1. All CD patients exhibited normal values for height, body weight, hemoglobin, AST, ALT, GFR, and creatinine levels. Acute phase protein (CRP) levels were >5 mg/dL in all patients except one (4 mg/dL). This outlier tested negative for stool calprotectin using the Simple Calprotectin one-step immunochromatographic test kit (Operon S.A., Spain), while the remaining CD patients were positive for stool calprotectin.

The healthy group (n=6; H1–H6) consisted of four males and two females aged 18–40 years. All healthy individuals were negative for stool calprotectin. Inclusion criteria for both CD patients and healthy individuals are provided in Table S1.

Given the exploratory nature of this study and the *in vitro* design, a sample size (<10 donors) is appropriate to capture biologically meaningful trends. Unlike large-scale clinical studies required to account for inter-individual variability, *in vitro* conditions provide a controlled environment that reduces this variability and allows us to investigate specific metabolic processes. Previous studies with similar sample sizes have successfully identified robust metabolic signatures and reproducible trends ^35,36,40^. Although our findings are supported by robust statistical analyses, future studies with larger cohorts will be necessary to validate and expand these results.

The study protocol was rigorously reviewed and approved by the Bioethics Committee of the University of Rzeszow (No. 2022/067) and adheres to the highest ethical standards in medical research.

### 2.2. In vitro batch culture experiments

*In vitro* tests of the metabolic activity of stool microbiotas were conducted in static batch cultures under anaerobic conditions (Figure 1). The liquid growth medium was a modified M9 ^41^ without CaCl_2_ and MgSO_4_, supplemented with 2 g/L yeast extract and (i) 10 g/L glucose (Becton, Dickinson and Company Sparks, USA) or (ii) 3.5 g/L sodium acetate (Chempur, Poland) and 7.41 g/L sodium lactate (VWR International). The growth media composition was inspired by our previous work ^42,43^. Stool samples were frozen immediately after collection from patients and healthy volunteers at −80 °C and stored under these conditions for one month prior to experimentation, without the addition of a cryoprotectant. To initiate each culture, 1 gram stool samples were added to 200 mL of the respective sterile media in 250-mL Erlenmeyer flasks and these were incubated for three days without shaking at 32°C in a Vinyl Anaerobic Chamber (Coy Laboratory Products, Inc., USA). The same volume of modified M9 medium without the added carbon substrates was also inoculated with the stool samples to characterize metabolites not resulting from the fermentation of glucose or acetate+lactate. In all cases the starting pH was 7.0 and no additional means of pH control were used. After three days of incubation, the cultures were centrifuged (10,000 × g for 10 min, 4°C) to remove microbial cells and debris. The supernatants were collected for chemical analysis of the post-fermentation liquids, while pellets were used for total microbial DNA isolation. Three independent cultures were performed for each stool sample in the three media: M9-based, M9-based plus glucose, M9-based plus acetate and lactate.

**Figure 1.**
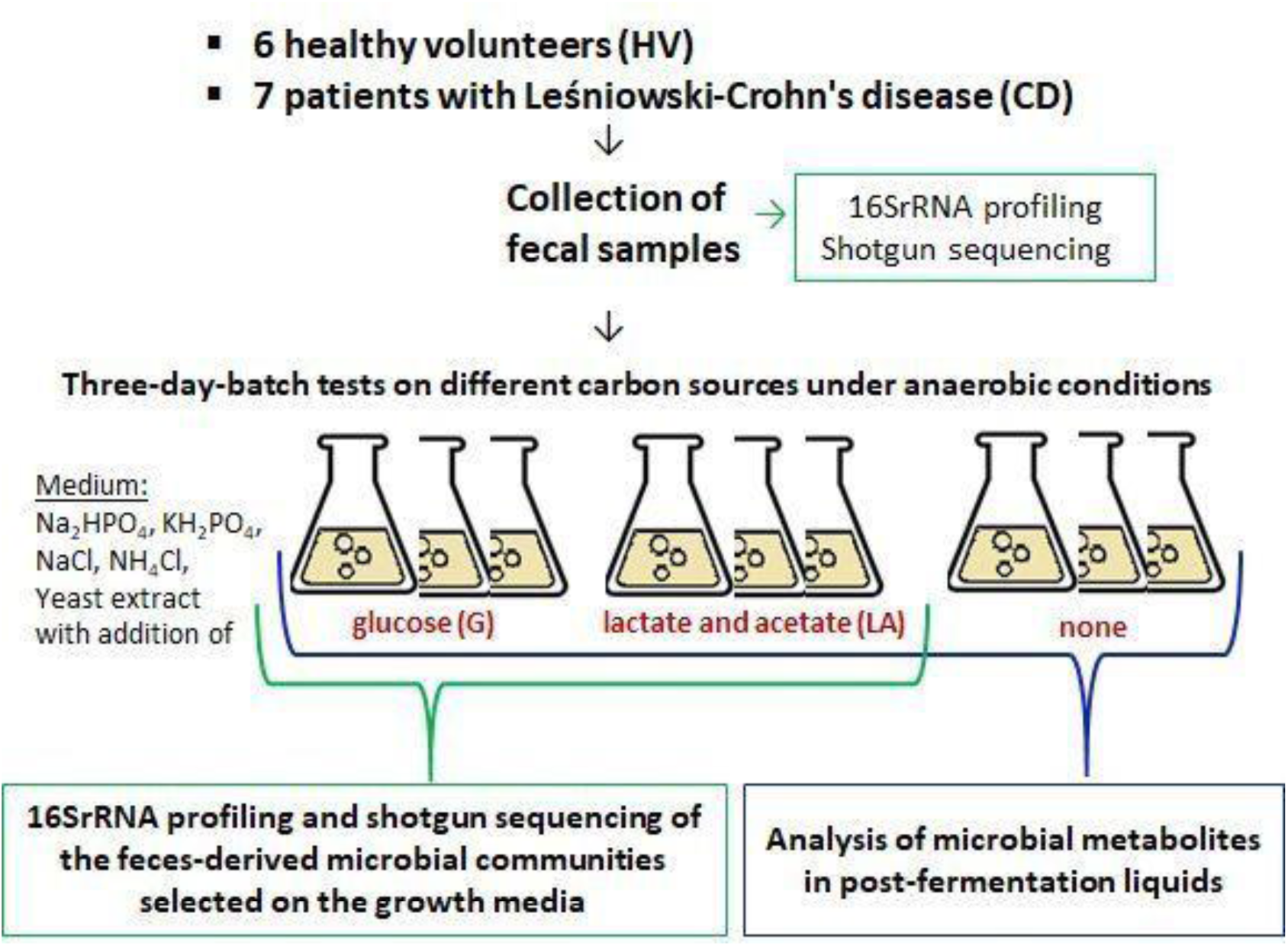
Scheme of the experimental setup.

### 2.3. Analytical methods

A standard pH meter (ELMETRON, Poland) was used to measure the pH of the media and post-fermentation liquids. Ethanol quantification in post-fermentation liquids was performed using a GC-MS method based on the protocol of Pinu et al.^44^. Derivatization of γ-aminobutyrate (GABA), L-alanine, L-leucine, L-isoleucine, and glycine followed the procedure outlined by Sun et al.^45^ (File S1). Chromatographic separations were carried out using DB-1701 and TG-5SILMS columns. Metabolite concentrations were determined via GC-MS using a Trace 1310 GC system coupled with a TSQ9000 Triple Quadrupole mass spectrometer (Thermo Scientific). Chromeleon 7 software was employed for instrument control and data acquisition, with MS data collected in Selected Reaction Monitoring mode. The analysis of C2–C6 organic acids, including acetic, lactic, propionic, butyric, 2-methyl butyric, isobutyric, isovaleric, valeric, isocaproic, and caproic acids, was performed as previously described ^46^. Calibration curves and internal standards were utilized for all quantified compounds.

### 2.4. Microbial DNA extraction

DNA was isolated from 0.2 g samples of stool and 0.2 g of material pelleted by centrifugation of 50 mL liquid cultures, using a QIAamp PowerFecal Pro DNA Kit (Qiagen, Cat. No. 51804) according to the manufacturer’s protocol. Cell lysis was achieved using a Vortex-Genie 2 equipped with a Vortex Adapter for 1.5-2 mL tubes (Scientific Industries, Inc., USA). DNA was eluted in 10 mM Tris-HCl (pH 8.5) and stored at −20°C until it was sequenced. Extraction blanks were included for contamination control.

### 2.5. 16S rRNA gene sequencing and bioinformatics

The V4 hypervariable region of the 16S rRNA gene was PCR-amplified from the prepared microbial DNA samples using the primer pair 515-F and 806-R ^47^, which carried Illumina adapters and a unique 12-base pair (bp) barcode identifying each sample. The PCR products were quantified with a Quant-iT PicoGreen dsDNA assay Kit (Invitrogen). The DNA fragments were then pooled in equimolar concentrations, and products longer than 200 bp were selected using QIAseq beads and sequenced on a 2×150 bp Illumina MiSeq platform (Illumina). Demultiplexing was performed using idemp (https://github.com/yhwu/idemp). The DADA2 pipeline was used to infer amplicon sequence variants (ASVs). The reads were trimmed to 145 bp, low-quality reads exceeding a maximum expected error of 2 bp were removed, and the remaining reads were used for error model training and correction. Reads were then merged, chimera sequences eliminated, and taxonomic identities assigned using the Ribosomal Database Project (RDP) classifier ^48^ with the SILVA nr version 138.1 rRNA database ^49^. The number of reads per sample after quality filtering, error correction, chimera removal, and taxonomy cleaning ranged from 50,154 to 97,882. The raw 16S rRNA DNA sequences have been deposited in NCBI databases with the accession number PRJNA1160194.

### 2.6. Shotgun metagenomics sequencing and bioinformatics

Stool culture samples from ten subjects (five healthy and five with CD) were randomly selected for shotgun metagenomic analysis. From the set of three biological replicates per substrate from each individual, the one closest to the group centroid calculated with Bray Curtis (BC) dissimilarities was sequenced. Additionally, an initial stool sample was sequenced for each individual patient. Quality filtering and adapter removal were performed with trimmomatic v 0.38 (LEADING:20 TRAILING:20 SLIDINGWINDOW:4:15 MINLEN:50) ^50^. Host contamination was removed following alignment to the human reference genome *GRCh38* using bowtie2 ^51^ with standard parameters, leaving between 23,651,272 to 62,620,933 read pairs per sample. Reads were taxonomically profiled using MetaPhlAn4 with the “mpa_vOct22_CHOCOPhlAnSGB_202212” database ^52^. The results were then imported into HUMAnN3 for functional profiling ^53^. The raw shotgun sequencing data have been deposited in NCBI databases with the accession number PRJNA1160072.

### 2.7. Statistical analysis

Statistical analyses were performed in R 4.2.1 ^54^ with the aid of the vegan package ^55^. Technical replicates were merged and considered as one sample. Alpha (ASV richness and Shannon’s diversity index) and beta diversity metrics were calculated on rarefied data (50,000 reads). Differences in alpha diversity were calculated with a linear mixed effects model with healthy stool samples as reference, subjects as random effects, and interactions between substrate and diagnosis. For beta diversity, BC dissimilarities were calculated on rarefied data ^56^. The effect of different groups on community dissimilarities was first visualized with non-metric multidimensional scaling (NMDS) plots and quantified through a Permutational multivariate analysis of variance (PERMANOVA) ^54^. Differential abundance of species and metabolites between combinations was also calculated with linear mixed effects models on log2 transformed relative abundances using the subject as a random effect, as implemented in the package maaslin2 ^57^.

## 3. Results

### 3.1. Fermentation products as a reflection of the metabolic activity of stool microbiota-derived communities

To evaluate whether the metabolic activity of stool microbiota *in vitro* corresponds to the condition of the gut microbiome *in vivo*, we analyzed fermentation product concentrations in post-fermentation liquids. Significant differences in metabolite levels (butyrate, isobutyrate, caproate, valerate, lactate, propionate, acetate, ethanol, γ-aminobutyrate, and amino acids) correlated with stool donor health status and the carbon source provided. To exclude metabolism derived from the inoculum, control cultures using M9 medium without additional carbon sources were assessed (Figure 1). These findings indicate that microbial metabolism follows distinct trajectories depending on the health status of the donor and the availability of specific carbon substrates. Detailed metabolite concentrations are available in Table S2.

Glucose fermentation led to a pH decrease from 7 to 4-5 (Figure 2a), while acetate and lactate metabolism maintained pH near neutrality, consistent with previous studies on microbial interactions and metabolic activity in anaerobic environments, including dark fermentation microbial communities ^42,43^. Principal Component Analysis (PCA) highlighted metabolite differences, with stool-only controls showing lower metabolite abundance than carbon-supplemented cultures (Figures 2b, 2c). Even in the absence of added carbon, CD-derived samples displayed elevated lactate levels and reduced C2-C6 organic acids, with valerate showing statistically significant differences (Figure 2d). Samples formed a metabolic gradient, with healthy microbiotas grown on acetate+lactate displaying higher C2-C6 organic acid production, whereas CD-associated communities grown on glucose exhibited increased ethanol, lactate, GABA, and amino acid levels (Figures 2b, 2c, 2d). These results suggest a fundamental shift in metabolic output between health-associated and CD-associated microbiotas.

**Figure 2.**
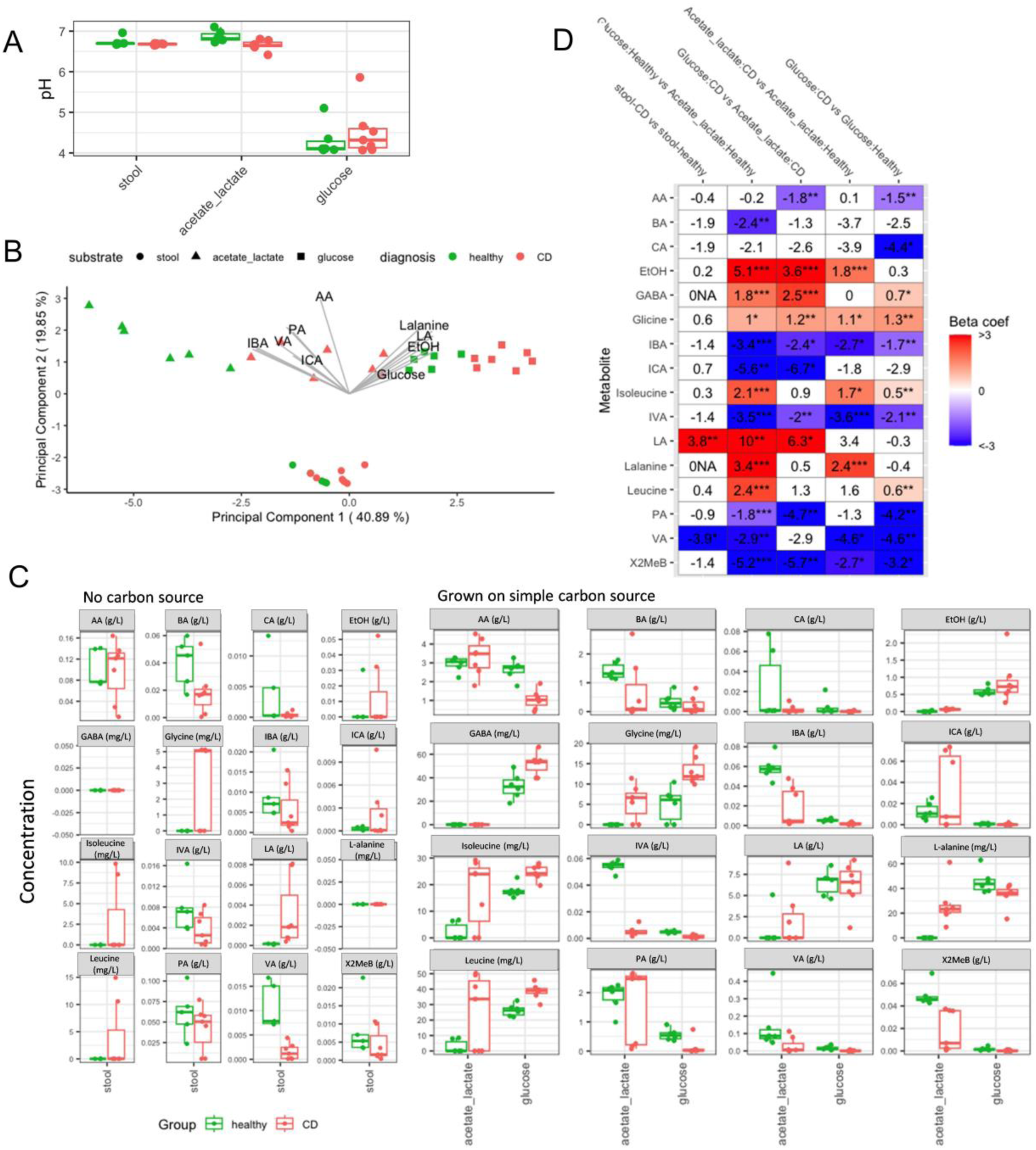
Outcomes of microbial metabolic activity in batch tests. (A) Boxplot showing the pH of post-fermentation liquids, green data points – healthy individuals, red data points – CD patients. (B) PCA ordination of metabolite measurements. Round points indicate baseline samples (samples without added carbon source). (C) Boxplots showing differences in the concentrations of metabolites across different groups. (D) Differential abundance analysis of metabolite concentrations. Beta-coefficient is equivalent to log2(fold change), P * <0.05, **<0.01, ***<0.001. Red cells indicate a significant increase, blue cells a significant decrease (P * <0.05, **<0.01, ***<0.001). White cells indicate non-significant differences. AA – acetate, BA – butyrate, CA – caproate, EtOH – ethanol, GABA – aminobutyrate, IBA – isobutyrate, ICA – isocaproate, IVA – isovalerate, LA – lactate, PA – propionate, VA – valerate, X2MeB – 2-methyl butyrate.

### 3.2. Microbial diversity in stool and in vitro cultures based on 16S rRNA gene sequencing

The analysis identified 1,662 distinct 16S rRNA gene amplicon sequence variants (ASVs). Linear mixed-effects models revealed a significant reduction in ASV richness and Shannon’s diversity (Shannon H’) in Crohn’s disease (CD) samples compared to healthy controls (Richness: β = -88.26, SE = 21.13, p < 0.001; Shannon H’: β = -0.92, SE = 0.28, p < 0.001; Figure 3a). This aligns with the well-documented reduction in gut microbiome diversity observed in CD patients. The decline in ASV richness and Shannon H’ was consistent across stool samples and *in vitro* cultivated microbial communities. Additionally, microbial communities selected on glucose exhibited lower diversity than those in raw stool samples (Richness: β = -58.41, SE = 15.99, p < 0.001; Shannon H’: β = -1.31, SE = 0.28, p < 0.001; Figure 3a). In contrast, no significant differences were detected in communities cultivated on acetate+lactate-containing medium. Detailed model outputs are provided in Tables S3 and S4.

**Figure 3.**
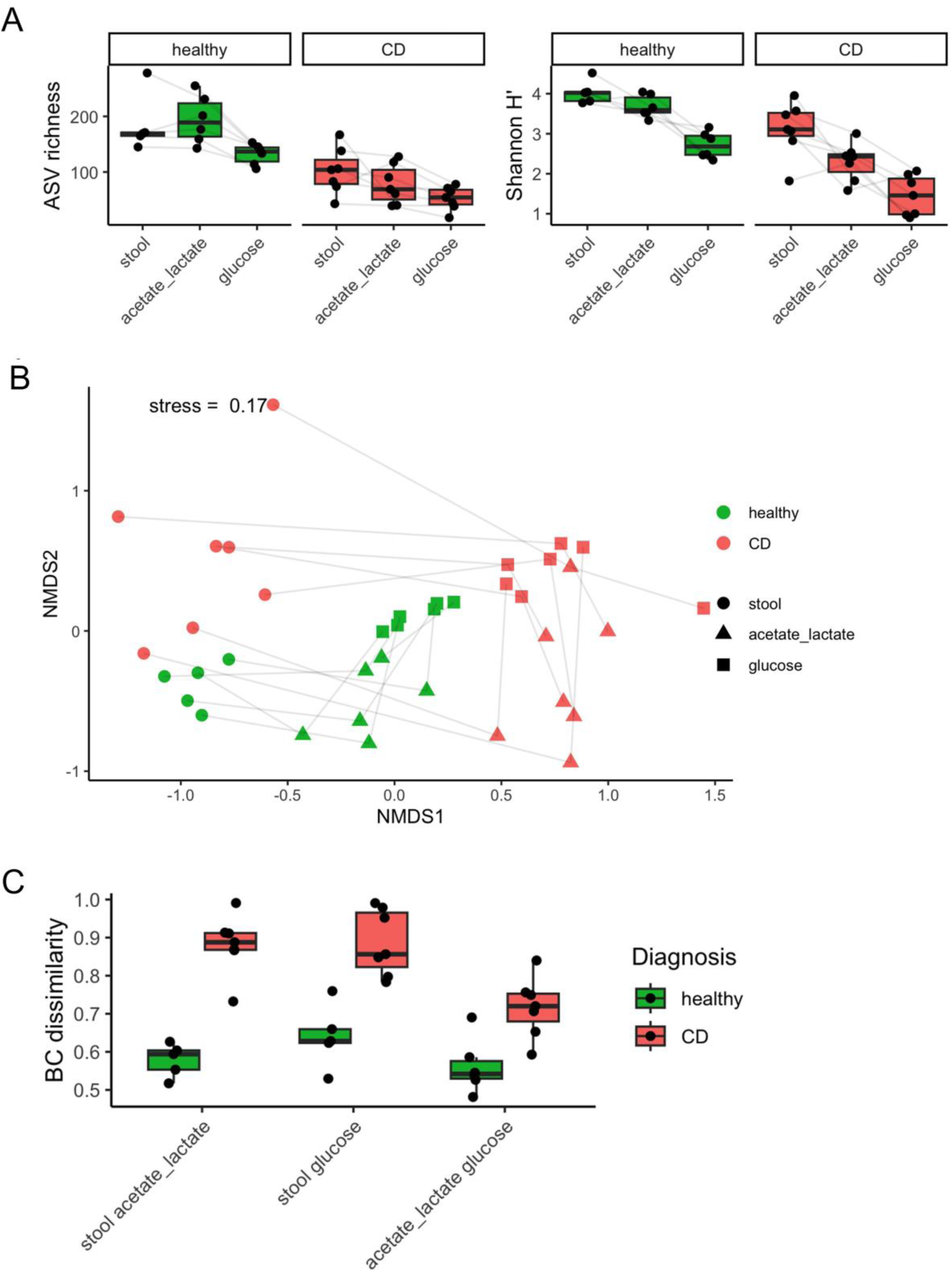
Results of 16S rRNA gene sequencing, illustrating biodiversity of the stool microbiota and stool-derived microbial communities selected by glucose or acetate+lactate. (A) Alpha diversity metrics. (B) NMDS plot based on BC dissimilarities between samples. A gray line connects samples from the same donor. (C) Comparison of BC dissimilarities between samples from the same donor (on the X axis, dissimilarities between stool and acetate+lactate samples, stool and glucose samples, and acetate+lactate and glucose samples).

Bray-Curtis dissimilarity analysis identified carbon substrate as the primary driver of community variation, followed by donor health status (Figure 3b, Table 1). CD-derived communities exhibited greater dissimilarity across substrates than healthy microbiotas, suggesting metabolic instability in CD-associated microbial communities (Figure 3c).

**Table 1.**
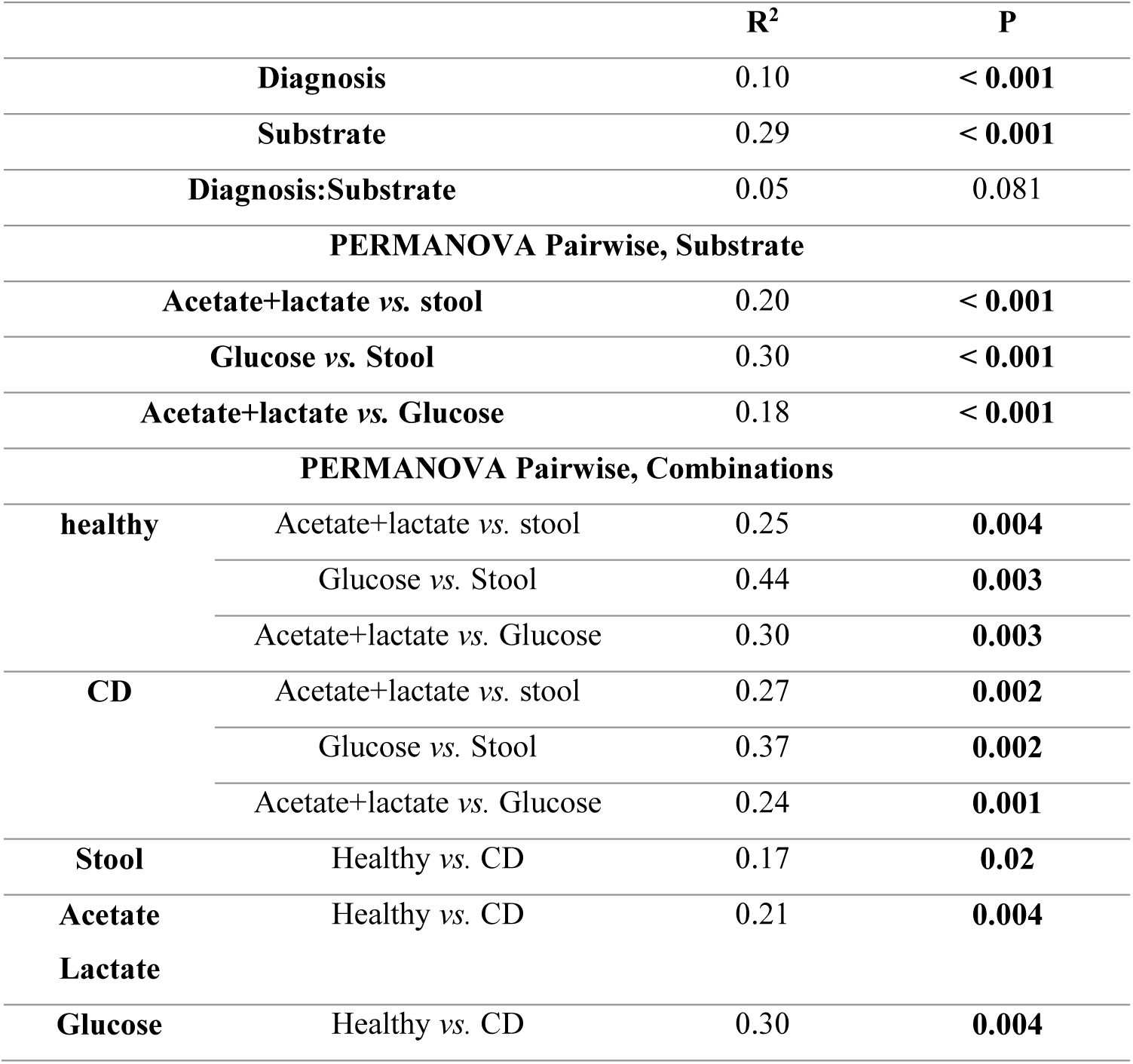
PERMANOVA statistics.

### 3.3. Metagenomic profiling of microbial communities in CD and healthy individuals

We performed metagenomic analysis on samples selected based on BC dissimilarities to investigate the CD-associated compositional shifts in microbial communities. From the three biological replicates per substrate, we chose the sample closest to the centroid for shotgun sequencing (see Methods). Taxonomic profiling using MetaPhlAn4 successfully assigned between 65% and 87% of the total taxonomic composition (Table S5). Batch culture experiments revealed a transition from highly diverse, low-abundance species to the dominance of specific taxa. Notably, microbial communities grown on glucose-enriched media exhibited an increased contribution of *Escherichia coli*, whereas acetate+lactate favored the proliferation of *Veillonella* spp. (Figure 4a). These shifts were particularly pronounced in CD-derived samples, suggesting a disease-specific microbial selection under distinct metabolic conditions. Differential abundance analysis confirmed these trends, highlighting additional taxa whose prevalence was substrate-dependent (Figure 4b, 4c).

**Figure 4.**
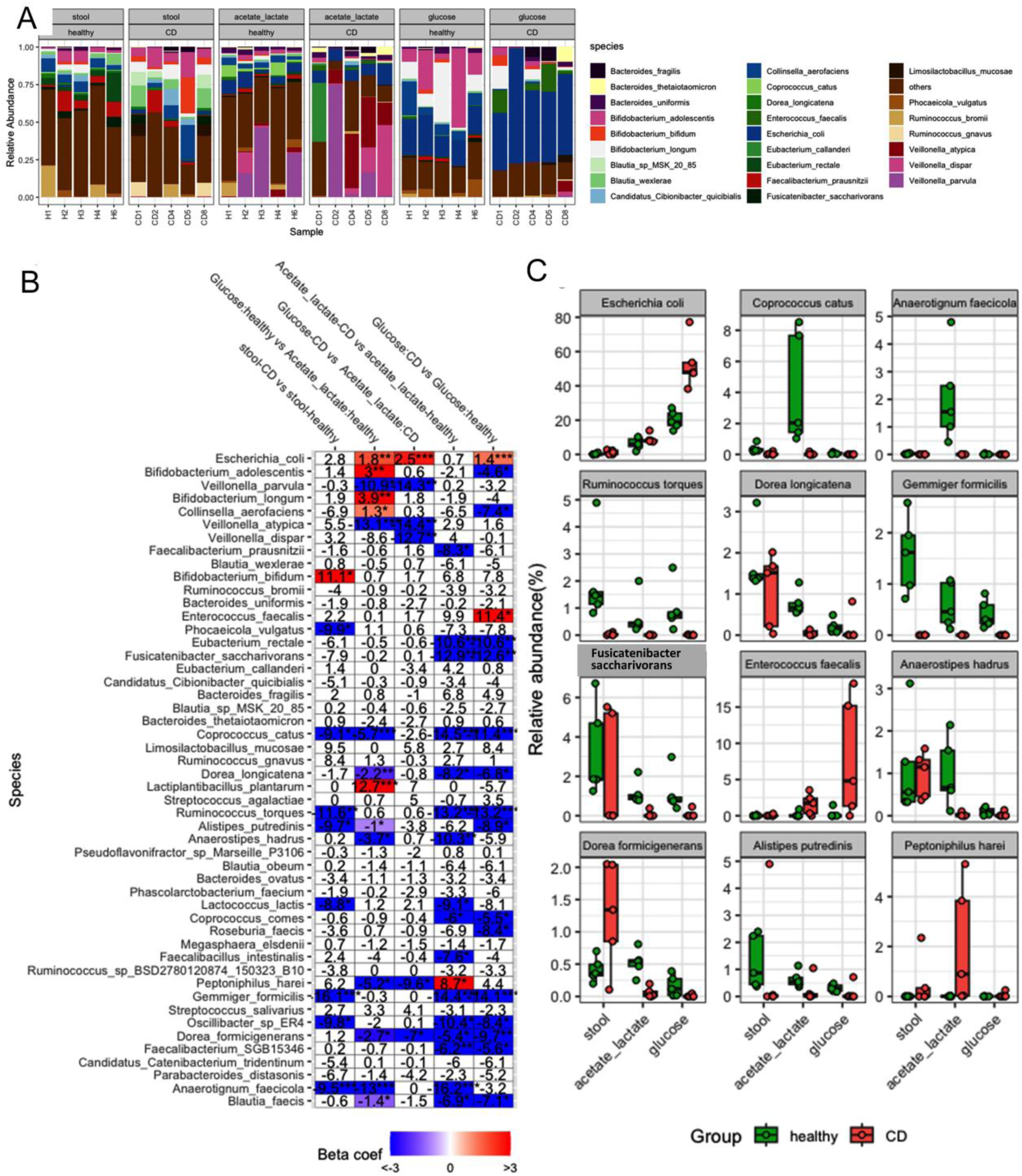
Taxonomic assignment of shotgun metagenomics reads. (A) Stacked bar plot of species composition as identified by MetaPhlAn4. Only the top 25 species across the study are presented; all others are grouped into “others.” (B) Differential abundance analysis of bacterial species. Beta-coefficient is equivalent to log2(fold change), P * <0.05, **<0.01, ***<0.001. Red cells indicate a significant increase, blue cells a significant decrease (P * <0.05, **<0.01, ***<0.001). White cells indicate non-significant differences. (C) Boxplots showing relative abundances across treatments of species selected as examples.

Comparison of baseline stool microbiota between CD patients and healthy individuals revealed a striking enrichment of *Bifidobacterium bifidum* in CD samples. However, the overarching trend was a marked reduction in species diversity in CD patients, with notable declines in e.g. *Phocaeicola vulgatus* and *Coprococcus catus* (Figure 4b, Table S6). This pattern persisted upon *in vitro* cultivation, with a significant decrease in species richness in CD-derived cultures compared to healthy controls across both glucose-and acetate+lactate-supplemented media. Some taxa, such as *Fusicatenibacter saccharivorans* and *Coprococcus catus*, exhibited reduced abundance in both media, whereas *Faecalibacterium prausnitzii* showed a selective reduction under acetate+lactate conditions (Table S6). Interestingly, *E. coli* and *Enterococcus faecalis* were significantly enriched in glucose-supplemented cultures from CD samples, mirroring their non-significant but elevated baseline levels in CD stool microbiota. Conversely, acetate+lactate supplementation promoted the growth of *Peptoniphilus harei* in CD-derived cultures.

Further comparison of microbial community responses in healthy versus CD-derived samples suggested a characteristic metabolic adaptability in healthy microbiota. In glucose-supplemented cultures, healthy samples exhibited preferential growth of *Bifidobacterium adolescentis, B. longum, Lactiplantibacillus plantarum,* and *Collinsella aerofaciens*, with *Bifidobacterium* sp. displaying a trend toward reduced abundance in CD cultures. Across both CD and healthy groups, acetate+lactate consistently promoted the proliferation of *Veillonella* spp., including *V. parvula* and *V. atypica*, suggesting a substrate-driven effect independent of health status.

These observed shifts in microbial composition may result from interspecies interactions and alterations in the chemical microenvironment induced by substrate fermentation. Future studies should explore the mechanistic basis of these changes, particularly their implications for CD-associated dysbiosis and disease progression.

### 3.4. Associations between bacterial species and metabolites

Associations between bacterial taxa and metabolite production, derived from Figures 2 and 4, are summarized in Figure 5. The analysis revealed that *E. coli* and *Lactiplantibacillus plantarum* were major contributors to the increased production of ethanol, lactate, GABA, and amino acids. This metabolic activity was particularly pronounced in microbial communities derived from CD patients, especially under glucose-rich conditions.

**Figure 5.**
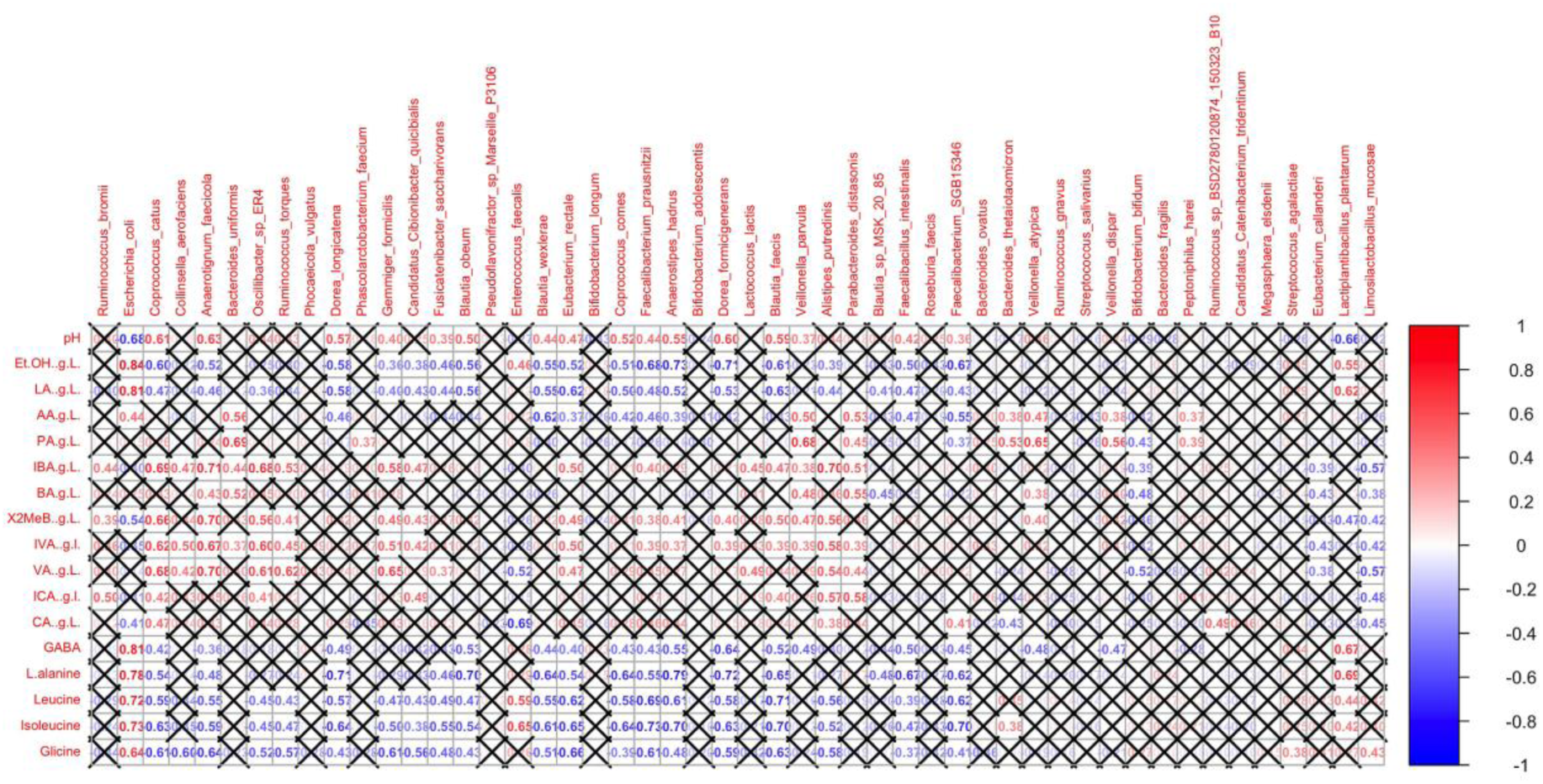
Spearman correlation matrix based on stool-derived microbial communities’ data showing correlations between bacterial species and the measured metabolites in *in vitro* batch cultures. Red: positive correlation, blue: negative correlation. Non-significant correlations (P < 0.05 after FDR correction) are marked with an ‘X’.

Conversely, species associated with higher levels of butyrate, valerate, and their derivatives exhibited an inverse correlation with ethanol, lactate, GABA, and amino acid synthesis. These species included *Coprococcus catus, Anaerotignum faecicola, Gemmiger formicilis, Eubacterium rectale, Alistipes putredinis, Blautia faecis*, and *Anaerostipes hadrus*. This pattern was most evident in microbial communities from healthy individuals cultivated in acetate+lactate-supplemented media. Interestingly, some species, such as *Parabacteroides distasonis* and *Bacteroides uniformis*, were associated with increased butyrate and valerate production without a corresponding decrease in other metabolites. In contrast, *Limosilactobacillus mucosae* and *Bifidobacterium bifidum* were linked to reduced levels of these compounds.

Elevated propionate production was strongly associated with *Phascolarctobacterium faecium,* while *Peptoniphilus harei* and *Veillonella dispar* correlated with both propionate and acetate synthesis. The production of caproate and iso-caproate frequently co-occurred with butyrate and valerate synthesis, though some species, such as *Ruminococcus*, exhibited a unique association with caproate production.

### 3.5. Metabolic potential analysis

To assess metabolic potential (Figure 6) using the HUMAnN3 tool, we selected genes encoding enzymes involved in reactions where the measured metabolites served as either products or substrates. Key fermentation pathways—including glycolysis, acetyl-CoA formation, hydrogen production, and the reductive carbon monoxide dehydrogenase/acetyl-CoA synthase (CODH/ACS) pathway—were also analyzed. A comprehensive list of enzymes and reactions under batch culture conditions is provided in Table S7.

**Figure 6.**
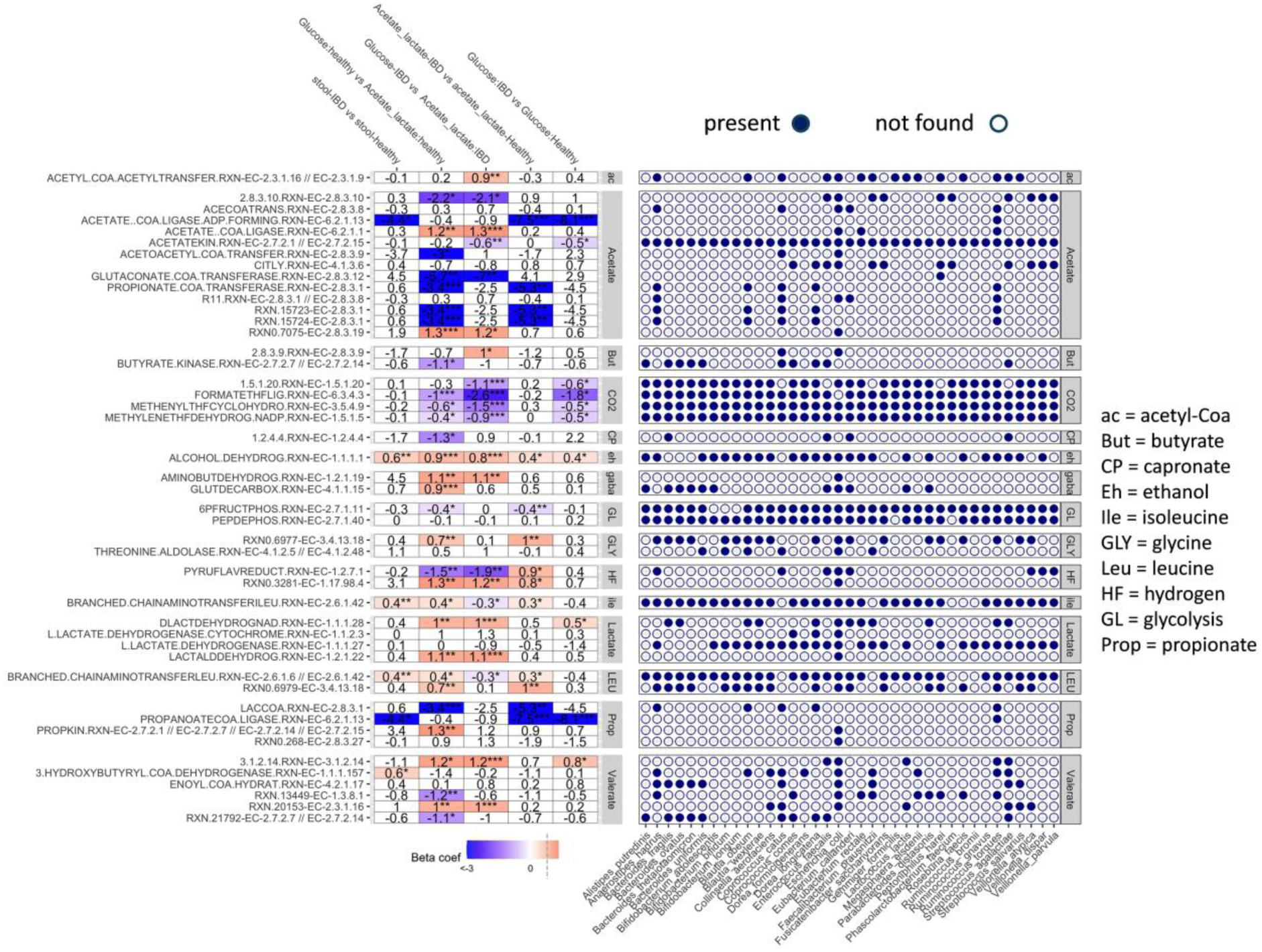
Differential abundance analysis of selected metabolites and fermentation processes and enzymes catalyzing reactions associated with them. Beta-coefficient is equivalent to log2(fold change), P * <0.05, **<0.01, ***<0.001. Red cells indicate a significant increase, blue cells a significant decrease (P * <0.05, **<0.01, ***<0.001). White cells indicate non-significant differences. The right-hand panel shows which of the genes encoding the different enzymes are present in selected species.

Stool samples from CD patients, compared to those from healthy individuals, exhibited an overrepresentation of genes encoding ethanol dehydrogenase (EC 1.1.1.1) and branched-chain amino acid transaminase (EC 2.6.1.42), both involved in leucine and isoleucine metabolism. Consequently, ethanol and amino acid overproduction by CD-derived microbial communities in *in vitro* cultures correlated with the upregulation of ethanol dehydrogenase (in both glucose- and acetate+lactate-supplemented media), dipeptidase (EC 3.4.13.18), and branched-chain amino acid transaminase (in acetate+lactate-enriched media).

CD stool-derived communities grown on acetate+lactate exhibited an increased hydrogen-producing potential via pyruvate:ferredoxin oxidoreductase (EC 1.2.7.1) and formate dehydrogenase (EC 1.17.98.4), consistent with clinical observations of bloating and gas production in IBD patients.

Distinct metabolic responses emerged in glucose-supplemented cultures. In healthy stool-derived cultures, glucose exposure reduced the contribution of genes encoding propionate CoA-transferase (EC 2.8.3.1), butyrate-acetoacetate CoA-transferase (EC 2.8.3.9), and 2-oxoisocaproate dehydrogenase (EC 1.2.4.4). By contrast, CD-associated microbiota exhibited increased expression of butyrate-acetoacetate CoA-transferase and acetyl-CoA C-acetyltransferase (EC 2.3.1.9), suggesting a metabolic shift favoring butyrate synthesis from glucose rather than acetate+lactate. Notably, a hallmark of a healthy microbiota is the efficient conversion of lactate and acetate into butyrate—a process that appears to be disrupted in CD-associated microbial communities.

Among all analyzed taxa, *Escherichia coli* emerged as the most metabolically versatile species. It dominated *in vitro* cultures supplemented with glucose, particularly in CD-derived communities, highlighting its ability to exploit diverse metabolic pathways. This metabolic flexibility may contribute to its persistence and overrepresentation in CD microbiota, reinforcing its role as a key player in disease-associated dysbiosis.

Genes encoding acetate kinase, lactate dehydrogenase (EC 1.1.1.27), glycolytic enzymes, and those involved in the reductive CODH/ACS pathway were consistently present across dominant taxa. However, genes associated with butyrate, propionate, and caproate synthesis were restricted to a subset of bacterial species and were absent in some taxa known to produce these metabolites. These unexpected findings underscore the need for further research and improved analytical methodologies to accurately capture metabolic potential.

Together, these findings illustrate the metabolic divergence between CD- and healthy-derived microbiotas, revealing substrate-driven microbial and functional shifts underlying CD-associated dysbiosis.

## 4. Discussion

The central hypothesis of this study was that differences in the composition, metabolic potential, and metabolic activities of CD-disrupted microbiomes can be effectively reflected in *in vitro* cultures of stool microbiota supplemented with simple organic carbon compounds found in the gut. These include sugars from dietary fiber degradation such as glucose and fermentation products such as acetate and lactate. We employed batch cultures using stool samples to inoculate media containing glucose or lactate with acetate. In this way we established stool microbiota conditions for (i) glycolytic fermentations, (ii) acetate and lactate conversion to butyrate ^37,38^, and (iii) propionate synthesis from lactate ^58^. The experimental setup ensured stable, controlled conditions, such as anaerobiosis, incubation time, and composition of bacterial growth media.

A key finding was that the fermentation pattern of fecal microbiota was altered in CD-patients compared to the healthy individuals due to disease-associated differences in microbial composition and function reproducible in batch cultures (Figure 7), offering indirect insights into the metabolic dynamics of gut microbiota.

**Figure 7.**
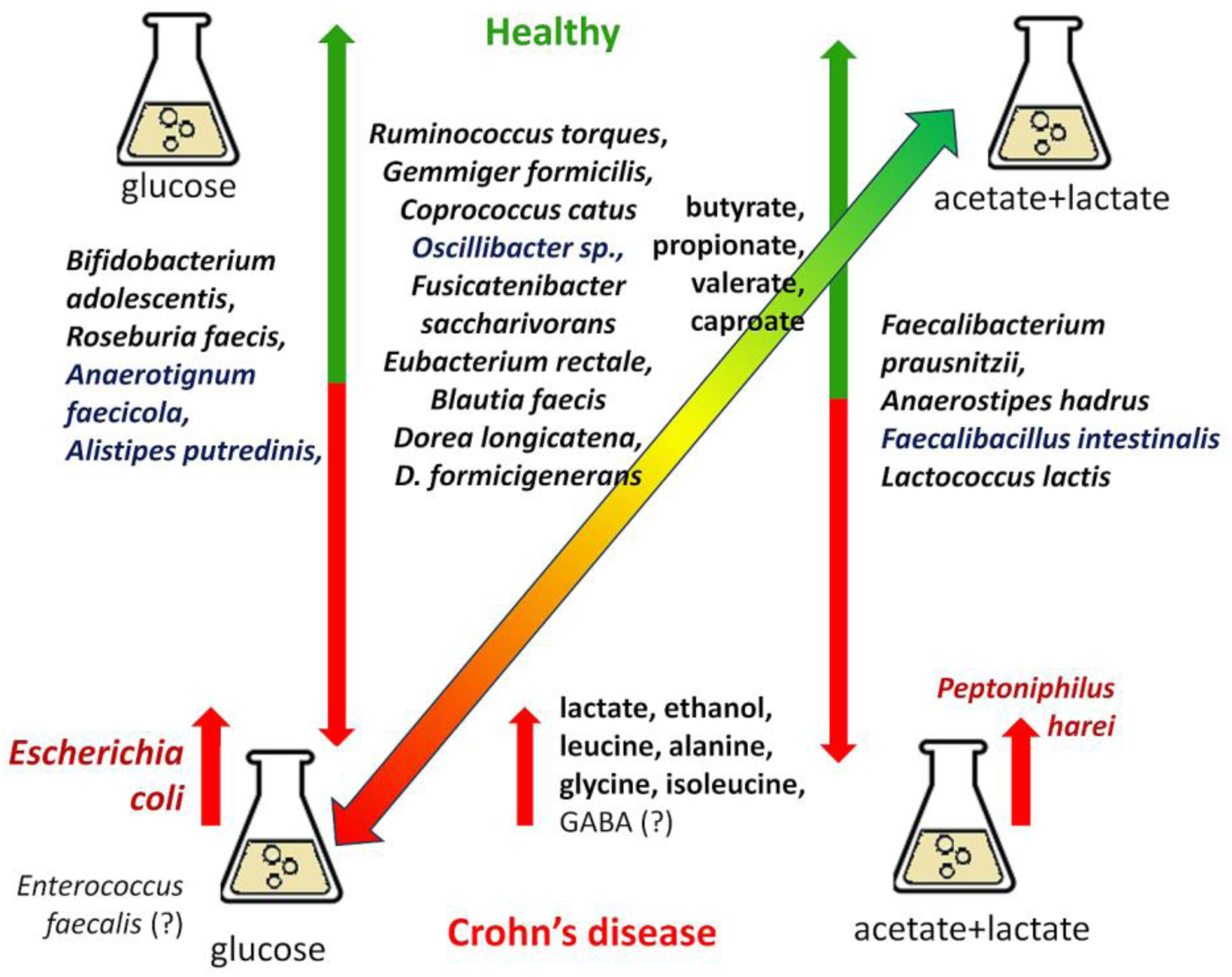
Summary of study findings: Crohn’s disease (CD)-associated dysbiosis features reduced synthesis of C2-C6 organic acids like butyrate, valerate, caproate, and propionate, coupled with elevated production of ethanol and amino acids. These metabolic shifts correspond to altered microbial composition, including depletion of beneficial commensals and overrepresentation of CD-associated bacteria, particularly *Escherichia coli*. *In vitro* fermentations reveal that acetate+lactate promote the growth of health-associated taxa, while glucose favors CD-linked microbes, highlighting distinct metabolic imbalances. Identifying key metabolites and microbial shifts may offer the potential for noninvasive diagnostics and even microbiome-targeted therapies in CD. Green indicates a healthy state, while red represents dysbiosis associated with Crohn’s disease. Downward arrows indicate a decrease and upward arrows signify an increase. Bacteria marked in blue are those whose role in the microbiome requires further investigation. Question marks denote bacterial species or metabolites whose association with a particular condition is surprising and warrants additional research.

### 4.1. Key Taxa Influenced by Carbon Sources

Analysis of stool samples from CD patients and healthy individuals confirmed the well-recognized decrease in microbial richness and diversity characteristic of CD-associated dysbiosis ^1,9,13,23–28^. Using the z-score measure, we showed that in CD-derived microbial communities, regardless of the carbon source in the media, the stool-associated genomes were reduced compared to those in healthy-derived consortia. Furthermore, in the healthy-derived consortia, genomes associated with acetate and lactate were present in higher abundance than those associated with glucose.

The study identified bacterial species that were significantly reduced in CD stool microbiotas and CD-derived microbial communities grown on specific carbon sources (details in Table S6). These species included key butyrate- and propionate-producing bacteria, including *Ruminococcus torques*, *Gemmiger formicilis*, *Eubacterium rectale* (*Agathobacter rectalis*), *Fusicatenibacter saccharivorans*, *Roseburia faecis* (*Agathobacter faecis*), *Faecalibacterium prausnitzii*, *Anaerostipes hadrus*, *Coprococcus catus* and non-butyrate producers such as *Phocaeicola vulgatus* (formerly *Bacteroides vulgatus*), *Lactococcus lactis, Dorea longicatena* and *D. formicigenerans*, *Blautia faecis, Bifidobacterium adolescentis* all of which are associated with maintaining gut homeostasis ^12,17,33^. The depletion of these beneficial bacteria was accompanied by an increased abundance of *Escherichia coli* and to a lesser extent, *Enterococcus faecalis*, particularly in glucose-supplemented cultures. *E. coli*, particularly adherent-invasive strains, is a well-known pathobiont linked to CD and other inflammatory conditions, with a capacity to utilize propionate and induce virulence ^59–61^. Our analysis of metabolic potential also revealed a propionate degradation pathway (PROPKIN-RXN) in *E. coli*. The role of *E. faecalis* in CD remains ambiguous given its dual capacity as both a probiotic and a potential driver of gut inflammation^62^. Its elevated level may reflect CD-related dysbiosis resulting from a decline in butyrate-producing bacteria and imbalances among key physiological bacterial groups. This dysregulation could also account for the observed higher prevalence of *Bifidobacterium bifidum* in the stools of CD patients compared to healthy individuals ^63^. Conflicting results regarding the *Bifidobacterium* species observed in this and other studies^1^ highlight the need for further research on species-specific effects within this genus.

Another notable observation was the stimulation of *Peptoniphilus harei* in CD microbiotas grown on acetate+lactate, which has not been extensively studied in the context of CD but is recognized as a commensal with opportunistic pathogenic potential ^64^. The reduced abundance of *Oscillibacter sp.*, *Anaerotignum faecicola*, *Alistipes putredinis*, and *Faecalibacillus intestinalis* (details in Table S6) in CD-related microbial communities suggests their potentially beneficial roles in the gut microbiome, warranting further investigation.

### 4.2. Metabolic activity in vitro and dysbiosis

Alterations in bacterial composition drive significant changes in the concentrations of fermentation products in *in vitro* stool microbiota cultures, enabling the reliable identification of metabolites whose levels may serve as potential metabolic biomarkers of health and disease. CD stool-derived microbial communities exhibited decreased production of propionate, butyrate, valerate, caproate, and their derivatives. These shifts align with the well-established role of butyrate and propionate in gut homeostasis and inflammation regulation ^1,3,9,13,23–28,65^. Importantly, our study highlights the inability of CD-derived microbiotas to effectively convert acetate and lactate into butyrate, a capability retained by healthy microbiotas ^37,38^. Analysis of the metabolic activity of stool microbiotas from six healthy individuals showed that butyrate was produced in all cases when acetate and lactate were used as carbon substrates. In contrast, among the seven CD patients, this occurred in only two cases (Table S2). Notably, one of these two patients had normal blood CRP levels and no detectable fecal calprotectin. Despite the small study group, the inability of CD-associated microbiotas to transform lactate and acetate into butyrate under in vitro conditions is a clear trend. The production of butyrate from acetate and lactate results from symbiotic interactions between lactic acid bacteria and butyrate producers, a process known as lactate cross-feeding ^37,38^. It is well established that many cross-feeding interactions are disrupted or diminished in dysbiosis associated with diseases, including IBD ^66^. Although less well characterized, valerate and caproate are believed to contribute to intestinal homeostasis, with reports suggesting that their levels decrease in IBD and increase following probiotic therapy ^65,67^.

CD stool-derived microbial communities exhibited altered amino acid metabolism, with increased alanine, glycine, leucine, and isoleucine concentrations in post-fermentation liquids from both cultures on glucose- and acetate+lactate-containing media. These amino acids, recognized as markers of CD-related dysbiosis ^9,23,31^ and are likely produced via pathways involving pyruvate and acetyl-CoA ^68,69^ from glucose, acetate, and lactate metabolism.

Ethanol and GABA were also notable metabolites produced by CD stool microbiotas grown *in vitro*, with ethanol emerging as a byproduct of acetate and lactate metabolism ^43^, and GABA being unexpectedly elevated in glucose cultures. While GABA production is typically reduced in IBD, its increase in batch cultures may result from microbiome dysregulation in CD, as previously described for *E. faecalis* and *B. bifidum* ^63,70^. We interpret this as a secondary effect of microbiome disturbances. Bacterial taxa with reduced abundance in CD stools and CD stool-derived microbial communities showed a positive correlation with butyrate, caproate, valerate, and their derivatives in post-fermentation liquids, while exhibiting a negative correlation with ethanol and amino acid levels. Notably, *E. coli* was positively correlated with all metabolic alterations characteristic of CD-derived communities.

### 4.3. Potential implications for diagnosis and therapy

Our findings suggest the utility of batch cultures of the stool microbiotas as a controlled system for reproducing CD-associated dysbiosis, enabling the identification of specific fermentation patterns and key metabolic products relevant to the disease. Their detection in post-fermentation supernatants, along with the identification of metabolic pathways—such as butyrate synthesis via lactate and acetate conversion— may provide a standardized, cost-effective, and non-invasive approach for assessing microbiome activity. Establishing reference concentrations for key diagnostic metabolites, including C2–C6 organic acids and amino acids, in larger cohorts might enhance dysbiosis detection and inform therapeutic strategies. Moreover, *in vitro* models offer a valuable platform for testing microbiome-targeted interventions, e.g. ^18,19^ aimed at restoring gut homeostasis in CD patients.

## 5. Conclusions

The dynamic and metabolic activity of stool microbiota grown in batch cultures with glucose or acetate and lactate enabled clear differentiation between CD-associated dysbiosis and a healthy gut microbiome. CD stool-derived microbial communities display lower microbial richness and diversity, and a loss of key C2-C6 organic acids producers. CD-associated microbiota cultivated in vitro show reduced production of valerate, caproate, and propionate and increased synthesis of ethanol and amino acids. The conversion of lactate and acetate to butyrate, a key function of a healthy gut microbiome, is significantly impaired in CD-associated microbial communities. Along with an increased potential for butyrate production from glucose, this suggests differences in butyrate synthesis pathways in the gut between healthy individuals and CD patients. *Escherichia coli* is a dominant species in CD-stool derived communities.. Its strong association with ethanol and amino acid production underscores the need for further research into its role in Crohn’s disease. The roles of *Peptoniphilus harei*, *Veillonella atypica, Enterococcus faecalis*, and *Bifidobacterium bifidum* in CD-associated dysbiosis remain unclear and warrant further study. These results may reinforce the potential value of batch culture systems for studying disease-related microbiome dynamics and suggest they could help identify specific metabolites that distinguish health from dysbiosis.

## Acknowledgments

This study was supported by This work was supported by (i) internal IBB PAS grant no. MG-2/21-18; (ii) the own sources of PANDA Core for Genomics and Microbiome Research, University of Arizona, and (iii) the own sources of the Institute of Medicine, Medical College of Rzeszow University.

## Author contributions

Anna Sikora, Anna Detman-Ignatowska planned the work and conceived and designed the experiments. Rafał Filip and Sara Jarmakiewicz-Czaja selected the study groups, obtained stool samples, and revised the manuscript. Anna Detman-Ignatowska, Anna Sikora, and Kinga Jakubowska performed the batch experiments. Anna Detman-Ignatowska isolated microbial DNA. Anna Williams, Nabahi Ramos Hickmanb, and Daniel Laubitz performed DNA sequencing. Gabriele Schiro analyzed metagenomic data, including data processing and proposing the conceptual framework for data interpretation and presentation. Emilia Samborowska and Jakub Karczmarski performed analyses of microbial metabolites. Anna Sikora, Gabriele Schiro, Anna Detman-Ignatowska wrote the manuscript. All authors read and approved the final manuscript.

## Supporting Information

**Table S1.** Clinical characteristics of the Crohn’s disease patients.

**Table S2.** Concentrations of metabolites (fermentation products) in post-fermentation liquids from batch cultures. Data used for the preparation of Figure 2 and Figure 5.

**Table S3.** Detailed results of linear mixed effects model (ASV richness). Data used for the preparation of Figure 3.

**Table S4.** Detailed results of linear mixed effects model (Shannon’s diversity). Data used for the preparation of Figure 3.

**Table S5.** Detailed taxonomic composition of the selected stool and stool-derived microbial communities grown on glucose- and acetate+lactate-containing media, based on shotgun metagenomic sequencing data.

**Table S6.** Bacterial species showing statistically significant changes in the fecal microbiota of patients with Crohn’s disease or in CD-stool-derived microbial communities. Table performed based on the data in Figure 4b. **Table S7.** Complete list of selected enzymes and biochemical reactions possible under the conditions applied in batch cultures, prepared using HUMAnN3. Data used for the preparation of Figure 6.

**File S1.** Detailed method of ethanol, GABA, alanine, leucine, isoleucine, and glycine determination.

## References

1. Pandey H., Jain D., Tang DWT., Wong SH., Lal D. Gut microbiota in pathophysiology, diagnosis, and therapeutics of inflammatory bowel disease. Intest Res 2024;22(1):15–43. Doi: 10.5217/ir.2023.00080.

2. The Human Microbiome Project Consortium Structure, function and diversity of the healthy human microbiome. Nature 2012;486(7402):207–14. Doi: 10.1038/nature11234.

3. Lloyd-Price J., Abu-Ali G., Huttenhower C. The healthy human microbiome. Genome Med 2016;8(1):51. Doi: 10.1186/s13073-016-0307-y.

4. Kundu P., Blacher E., Elinav E., Pettersson S. Our Gut Microbiome: The Evolving Inner Self. Cell 2017;171(7):1481–93. Doi: 10.1016/j.cell.2017.11.024.

5. Beura S., Kundu P., Das AK., Ghosh A. Metagenome-scale community metabolic modelling for understanding the role of gut microbiota in human health. Computers in Biology and Medicine 2022;149:105997. Doi: 10.1016/j.compbiomed.2022.105997.

6. Kuziel GA., Rakoff-Nahoum S. The gut microbiome. Current Biology 2022;32(6):R257–64. Doi: 10.1016/j.cub.2022.02.023.

7. Koh A., De Vadder F., Kovatcheva-Datchary P., Bäckhed F. From Dietary Fiber to Host Physiology: Short-Chain Fatty Acids as Key Bacterial Metabolites. Cell 2016;165(6):1332–45. Doi: 10.1016/j.cell.2016.05.041.

8. Rowland I., Gibson G., Heinken A., Scott K., Swann J., Thiele I., et al. Gut microbiota functions: metabolism of nutrients and other food components. Eur J Nutr 2018;57(1):1–24. Doi: 10.1007/s00394-017-1445-8.

9. Upadhyay KG., Desai DC., Ashavaid TF., Dherai AJ. Microbiome and metabolome in inflammatory bowel disease. J of Gastro and Hepatol 2023;38(1):34–43. Doi: 10.1111/jgh.16043.

10. Van Der Hee B., Wells JM. Microbial Regulation of Host Physiology by Short-chain Fatty Acids. Trends in Microbiology 2021;29(8):700–12. Doi: 10.1016/j.tim.2021.02.001.

11. Lloyd-Price J., Arze C., Ananthakrishnan AN., Schirmer M., Avila-Pacheco J., Poon TW., et al. Multi-omics of the gut microbial ecosystem in inflammatory bowel diseases. Nature 2019;569(7758):655–62. Doi: 10.1038/s41586-019-1237-9.

12. Ning L., Zhou Y-L., Sun H., Zhang Y., Shen C., Wang Z., et al. Microbiome and metabolome features in inflammatory bowel disease via multi-omics integration analyses across cohorts. Nat Commun 2023;14(1):7135. Doi: 10.1038/s41467-023-42788-0.

13. Madhogaria B., Bhowmik P., Kundu A. Correlation between human gut microbiome and diseases. Infectious Medicine 2022;1(3):180–91. Doi: 10.1016/j.imj.2022.08.004.

14. Young VB. The role of the microbiome in human health and disease: an introduction for clinicians. BMJ 2017:j831. Doi: 10.1136/bmj.j831.

15. Shaffer M., Armstrong AJS., Phelan VV., Reisdorph N., Lozupone CA. Microbiome and metabolome data integration provides insight into health and disease. Translational Research 2017;189:51–64. Doi: 10.1016/j.trsl.2017.07.001.

16. Lynch SV., Pedersen O. The Human Intestinal Microbiome in Health and Disease. N Engl J Med 2016;375(24):2369–79. Doi: 10.1056/NEJMra1600266.

17. Pittayanon R., Lau JT., Leontiadis GI., Tse F., Yuan Y., Surette M., et al. Differences in Gut Microbiota in Patients With vs Without Inflammatory Bowel Diseases: A Systematic Review. Gastroenterology 2020;158(4):930–946.e1. Doi: 10.1053/j.gastro.2019.11.294.

18. Ozbey D., Saribas S., Kocazeybek B. Gut microbiota in Crohn’s disease pathogenesis. World J Gastroenterol 2025;31(6). Doi: 10.3748/wjg.v31.i6.101266.

19. Zhang T., Li X., Li J., Sun F., Duan L. Gut microbiome-targeted therapies as adjuvant treatments in inflammatory bowel diseases: a systematic review and network meta-analysis. J of Gastro and Hepatol 2025;40(1):78–88. Doi: 10.1111/jgh.16795.

20. Buie MJ., Quan J., Windsor JW., Coward S., Hansen TM., King JA., et al. Global Hospitalization Trends for Crohn’s Disease and Ulcerative Colitis in the 21st Century: A Systematic Review With Temporal Analyses. Clinical Gastroenterology and Hepatology 2023;21(9):2211–21. Doi: 10.1016/j.cgh.2022.06.030.

21. Vich Vila A., Imhann F., Collij V., Jankipersadsing SA., Gurry T., Mujagic Z., et al. Gut microbiota composition and functional changes in inflammatory bowel disease and irritable bowel syndrome. Sci Transl Med 2018;10(472):eaap8914. Doi: 10.1126/scitranslmed.aap8914.

22. Bruner LP., White AM., Proksell S. Inflammatory Bowel Disease. Primary Care: Clinics in Office Practice 2023;50(3):411–27. Doi: 10.1016/j.pop.2023.03.009.

23. Mu C., Zhao Q., Zhao Q., Yang L., Pang X., Liu T., et al. Multi-omics in Crohn’s disease: New insights from inside. Computational and Structural Biotechnology Journal 2023;21:3054–72. Doi: 10.1016/j.csbj.2023.05.010.

24. Borren NZ., Plichta D., Joshi AD., Bonilla G., Peng V., Colizzo FP., et al. Alterations in Fecal Microbiomes and Serum Metabolomes of Fatigued Patients With Quiescent Inflammatory Bowel Diseases. Clinical Gastroenterology and Hepatology 2021;19(3):519–527.e5. Doi: 10.1016/j.cgh.2020.03.013.

25. Stange EF., Schroeder BO. Microbiota and mucosal defense in IBD: an update. Expert Review of Gastroenterology & Hepatology 2019;13(10):963–76. Doi: 10.1080/17474124.2019.1671822.

26. Lee M., Chang EB. Inflammatory Bowel Diseases (IBD) and the Microbiome—Searching the Crime Scene for Clues. Gastroenterology 2021;160(2):524–37. Doi: 10.1053/j.gastro.2020.09.056.

27. Glassner KL., Abraham BP., Quigley EMM. The microbiome and inflammatory bowel disease. Journal of Allergy and Clinical Immunology 2020;145(1):16–27. Doi: 10.1016/j.jaci.2019.11.003.

28. Khan I., Ullah N., Zha L., Bai Y., Khan A., Zhao T., et al. Alteration of Gut Microbiota in Inflammatory Bowel Disease (IBD): Cause or Consequence? IBD Treatment Targeting the Gut Microbiome. Pathogens 2019;8(3):126. Doi: 10.3390/pathogens8030126.

29. Deleu S., Machiels K., Raes J., Verbeke K., Vermeire S. Short chain fatty acids and its producing organisms: An overlooked therapy for IBD? eBioMedicine 2021;66:103293. Doi: 10.1016/j.ebiom.2021.103293.

30. Gallagher K., Catesson A., Griffin JL., Holmes E., Williams HRT. Metabolomic Analysis in Inflammatory Bowel Disease: A Systematic Review. Journal of Crohn’s and Colitis 2021;15(5):813–26. Doi: 10.1093/ecco-jcc/jjaa227.

31. Vich Vila A., Hu S., Andreu-Sánchez S., Collij V., Jansen BH., Augustijn HE., et al. Faecal metabolome and its determinants in inflammatory bowel disease. Gut 2023;72(8):1472–85. Doi: 10.1136/gutjnl-2022-328048.

32. Franzosa EA., Sirota-Madi A., Avila-Pacheco J., Fornelos N., Haiser HJ., Reinker S., et al. Gut microbiome structure and metabolic activity in inflammatory bowel disease. Nat Microbiol 2018;4(2):293–305. Doi: 10.1038/s41564-018-0306-4.

33. Gonzalez CG., Mills RH., Zhu Q., Sauceda C., Knight R., Dulai PS., et al. Location-specific signatures of Crohn’s disease at a multi-omics scale. Microbiome 2022;10(1):133. Doi: 10.1186/s40168-022-01331-x.

34. Biagini F., Daddi C., Calvigioni M., De Maria C., Zhang YS., Ghelardi E., et al. Designs and methodologies to recreate in vitro human gut microbiota models. Bio-Des Manuf 2023;6(3):298–318. Doi: 10.1007/s42242-022-00210-6.

35. Chen W., Tan D., Yang Z., Tang J., Bai W., Tian L. Fermentation patterns of prebiotics fructooligosaccharides-SCFA esters inoculated with fecal microbiota from ulcerative colitis patients. Food and Chemical Toxicology 2023;180:114009. Doi: 10.1016/j.fct.2023.114009.

36. Xu L., Yu Q., Ma L., Su T., Zhang D., Yao D., et al. In vitro simulated fecal fermentation of mixed grains on short-chain fatty acid generation and its metabolized mechanism. Food Research International 2023;170:112949. Doi: 10.1016/j.foodres.2023.112949.

37. Duncan SH., Barcenilla A., Stewart CS., Pryde SE., Flint HJ. Acetate Utilization and Butyryl Coenzyme A (CoA):Acetate-CoA Transferase in Butyrate-Producing Bacteria from the Human Large Intestine. Appl Environ Microbiol 2002;68(10):5186–90. Doi: 10.1128/AEM.68.10.5186-5190.2002.

38. Duncan SH., Louis P., Flint HJ. Lactate-Utilizing Bacteria, Isolated from Human Feces, That Produce Butyrate as a Major Fermentation Product. Appl Environ Microbiol 2004;70(10):5810–7. Doi: 10.1128/AEM.70.10.5810-5817.2004.

39. Tibble J. A simple method for assessing intestinal inflammation in Crohn’s disease. Gut 2000;47(4):506–13. Doi: 10.1136/gut.47.4.506.

40. Shehata E., Day-Walsh P., Kellingray L., Narbad A., Kroon PA. Spontaneous and Microbiota-Driven Degradation of Anthocyanins in an In Vitro Human Colon Model. Molecular Nutrition Food Res 2023;67(19):2300036. Doi: 10.1002/mnfr.202300036.

41. Miller JH. Experiments in Molecular Genetics, Cold Spring Harbor, NY: Cold Spring Harbor Laboratory Press; 1972.

42. Detman A., Laubitz D., Chojnacka A., Kiela PR., Salamon A., Barberán A., et al. Dynamics of dark fermentation microbial communities in the light of lactate and butyrate production. Microbiome 2021;9(1):158. Doi: 10.1186/s40168-021-01105-x.

43. Detman A., Mielecki D., Chojnacka A., Salamon A., Błaszczyk MK., Sikora A. Cell factories converting lactate and acetate to butyrate: Clostridium butyricum and microbial communities from dark fermentation bioreactors. Microb Cell Fact 2019;18(1):36. Doi: 10.1186/s12934-019-1085-1.

44. Pinu F., Villas-boas SG. Rapid Quantification of Major Volatile Metabolites in Fermented Food and Beverages Using Gas Chromatography-Mass Spectrometry. Metabolites 2017;7(3):37. Doi: 10.3390/metabo7030037.

45. Sun S., Wang H., Xie J., Su Y. Simultaneous determination of rhamnose, xylitol, arabitol, fructose, glucose, inositol, sucrose, maltose in jujube (Zizyphus jujube Mill.) extract: comparison of HPLC– ELSD, LC–ESI–MS/MS and GC–MS. Chemistry Central Journal 2016;10(1):25. Doi: 10.1186/s13065-016-0171-2.

46. Ostrowska J., Samborowska E., Jaworski M., Toczyłowska K., Szostak-Węgierek D. The Potential Role of SCFAs in Modulating Cardiometabolic Risk by Interacting with Adiposity Parameters and Diet. Nutrients 2024;16(2):266. Doi: 10.3390/nu16020266.

47. Caporaso JG., Lauber CL., Walters WA., Berg-Lyons D., Huntley J., Fierer N., et al. Ultra-high-throughput microbial community analysis on the Illumina HiSeq and MiSeq platforms. The ISME Journal 2012;6(8):1621–4. Doi: 10.1038/ismej.2012.8.

48. Wang Q., Garrity GM., Tiedje JM., Cole JR. Naïve Bayesian Classifier for Rapid Assignment of rRNA Sequences into the New Bacterial Taxonomy. Appl Environ Microbiol 2007;73(16):5261–7. Doi: 10.1128/AEM.00062-07.

49. Quast C., Pruesse E., Yilmaz P., Gerken J., Schweer T., Yarza P., et al. The SILVA ribosomal RNA gene database project: improved data processing and web-based tools. Nucleic Acids Research 2012;41(D1):D590–6. Doi: 10.1093/nar/gks1219.

50. Bolger AM., Lohse M., Usadel B. Trimmomatic: a flexible trimmer for Illumina sequence data. Bioinformatics 2014;30(15):2114–20. Doi: 10.1093/bioinformatics/btu170.

51. Langmead B., Salzberg SL. Fast gapped-read alignment with Bowtie 2. Nat Methods 2012;9(4):357–9. Doi: 10.1038/nmeth.1923.

52. Blanco-Míguez A., Beghini F., Cumbo F., McIver LJ., Thompson KN., Zolfo M., et al. Extending and improving metagenomic taxonomic profiling with uncharacterized species using MetaPhlAn 4. Nat Biotechnol 2023;41(11):1633–44. Doi: 10.1038/s41587-023-01688-w.

53. Beghini F., McIver LJ., Blanco-Míguez A., Dubois L., Asnicar F., Maharjan S., et al. Integrating taxonomic, functional, and strain-level profiling of diverse microbial communities with bioBakery 3. eLife 2021;10:e65088. Doi: 10.7554/eLife.65088.

54. Anderson MJ. A new method for non-parametric multivariate analysis of variance. Austral Ecology 2001;26(1):32–46. Doi: 10.1111/j.1442-9993.2001.01070.pp.x.

55. Oksanen J, Blanchet FG, Friendly M, Kindt R, Legendre P, McGlinn D, Minchin PR, O’Hara RB, Simpson GL, Solymos P, Stevens MHH, Szoecs E, Wagner H. vegan: Community Ecology Package. 2018.

56. Paulson JN., Stine OC., Bravo HC., Pop M. Differential abundance analysis for microbial marker-gene surveys. Nat Methods 2013;10(12):1200–2. Doi: 10.1038/nmeth.2658.

57. Mallick H., Rahnavard A., McIver LJ., Ma S., Zhang Y., Nguyen LH., et al. Multivariable association discovery in population-scale meta-omics studies. PLoS Comput Biol 2021;17(11):e1009442. Doi: 10.1371/journal.pcbi.1009442.

58. Reichardt N., Duncan SH., Young P., Belenguer A., McWilliam Leitch C., Scott KP., et al. Phylogenetic distribution of three pathways for propionate production within the human gut microbiota. The ISME Journal 2014;8(6):1323–35. Doi: 10.1038/ismej.2014.14.

59. Mirsepasi-Lauridsen HC., Vallance BA., Krogfelt KA., Petersen AM. *Escherichia coli* Pathobionts Associated with Inflammatory Bowel Disease. Clin Microbiol Rev 2019;32(2):e00060–18. Doi: 10.1128/CMR.00060-18.

60. Pobeguts OV., Ladygina VG., Evsyutina DV., Eremeev AV., Zubov AI., Matyushkina DS., et al. Propionate Induces Virulent Properties of Crohn’s Disease-Associated Escherichia coli. Front Microbiol 2020;11:1460. Doi: 10.3389/fmicb.2020.01460.

61. Ormsby MJ., Logan M., Johnson SA., McIntosh A., Fallata G., Papadopoulou R., et al. Inflammation associated ethanolamine facilitates infection by Crohn’s disease-linked adherent-invasive Escherichia coli. EBioMedicine 2019;43:325–32. Doi: 10.1016/j.ebiom.2019.03.071.

62. De Almeida CV., Taddei A., Amedei A. The controversial role of *Enterococcus faecalis* in colorectal cancer. Therap Adv Gastroenterol 2018;11:1756284818783606. Doi: 10.1177/1756284818783606.

63. Wang W., Chen L., Zhou R., Wang X., Song L., Huang S., et al. Increased Proportions of Bifidobacterium and the Lactobacillus Group and Loss of Butyrate-Producing Bacteria in Inflammatory Bowel Disease. J Clin Microbiol 2014;52(2):398–406. Doi: 10.1128/JCM.01500-13.

64. Wan X., Wang S., Wang M., Liu J., Zhang Y. Identification of Peptoniphilus harei From Blood Cultures in an Infected Aortic Aneurysm Patient: Case Report and Review Published Literature. Front Cell Infect Microbiol 2021;11:755225. Doi: 10.3389/fcimb.2021.755225.

65. Di Martino L., Osme A., Ghannoum M., Cominelli F. A Novel Probiotic Combination Ameliorates Crohn’s Disease–Like Ileitis by Increasing Short-Chain Fatty Acid Production and Modulating Essential Adaptive Immune Pathways. Inflammatory Bowel Diseases 2023;29(7):1105–17. Doi: 10.1093/ibd/izac284.

66. Marcelino VR., Welsh C., Diener C., Gulliver EL., Rutten EL., Young RB., et al. Disease-specific loss of microbial cross-feeding interactions in the human gut. Nat Commun 2023;14(1):6546. Doi: 10.1038/s41467-023-42112-w.

67. Markowiak-Kopeć P., Śliżewska K. The Effect of Probiotics on the Production of Short-Chain Fatty Acids by Human Intestinal Microbiome. Nutrients 2020;12(4):1107. Doi: 10.3390/nu12041107.

68. Wang Y., Xu J., Jin Z., Xia X., Zhang W. Improvement of acetyl-CoA supply and glucose utilization increases L -leucine production in *Corynebacterium glutamicum*. Biotechnology Journal 2022;17(8):2100349. Doi: 10.1002/biot.202100349.

69. Yamamoto S., Gunji W., Suzuki H., Toda H., Suda M., Jojima T., et al. Overexpression of Genes Encoding Glycolytic Enzymes in Corynebacterium glutamicum Enhances Glucose Metabolism and Alanine Production under Oxygen Deprivation Conditions. Appl Environ Microbiol 2012;78(12):4447–57. Doi: 10.1128/AEM.07998-11.

70. Strandwitz P., Kim KH., Terekhova D., Liu JK., Sharma A., Levering J., et al. GABA-modulating bacteria of the human gut microbiota. Nat Microbiol 2018;4(3):396–403. Doi: 10.1038/s41564-018-0307-3.

