## Supplementary material for "Dynamics and fermentation patterns of stool microbiota on simple carbon sources in *in vitro* batch cultures reveal dysbiosis in Crohn’s disease": File S1: File_S1.docx

Detailed method of ethanol, γ-aminobutyric acid (GABA), alanine, leucine, isoleucine, and glycine determination.

Ethanol was quantified using gas chromatography–mass spectrometry (GC/MS) on a TSQ9000 (Thermo Scientific) Triple Quadrupole mass spectrometer coupled to a Trace 1310 (Thermo Scientific) GC system, equipped with a DB-1701 column (30 m × 0.25 mm × 0.15 μm) (Agilent Technologies, CA, USA) and a TriPlus™ RSH Autosampler (Thermo Scientific). The injector temperature was set to 180°C, while the ion source, quadrupole, and transfer line temperatures were maintained at 250°C, 150°C, and 270°C, respectively. Helium was used as the carrier gas at a constant flow rate of 1 mL/min. A 1 μL aliquot of each sample was injected with a split ratio of 100:1. The column temperature was initially held at 50°C for 1 min, then increased at a rate of 20°C/min to 80°C, followed by a rapid ramp of 80°C/min to 270°C. Mass spectrometry data were collected in Selected Reaction Monitoring (SRM) mode. Target ions (m/z) for ethanol and d6-ethanol were used for the calibration curve and sample analysis.

Ethanol concentrations were determined using a calibration curve with an internal standard. Ethanol (calibration standard) and ethyl acetate were purchased from J.T.Baker (Phillipsburg, NJ, USA), while ethanol-d6 (labeled internal standard) was obtained from Sigma-Aldrich (St. Louis, MO, USA). All standard solutions were prepared using ultra-pure water (Milli-Q, Millipore, Milford, MA, USA). The GC-MS method for ethanol quantification in biological samples was adapted from ^1^.

For sample preparation, 400 µL of each sample was mixed with 10 µL of internal standard (7.9 mg/mL d6-ethanol) and 400 µL of ethyl acetate using an IKA® VRX basic Vibrax orbital shaker for 10 min. The samples were then centrifuged at 14,000 × g for 4 min, and 250 μL of the organic phase was transferred to GC-MS vials for analysis.

The concentrations of GABA, L-alanine, L-leucine, L-isoleucine and glycine were determined using a similar GC/MS approach, with the following modifications. A TG-5SILMS (Thermo) column was used for compound separation, and the injector temperature was set to 260°C. The initial column temperature was 70°C (held for 1 min), followed by a ramp of 20°C/min to 320°C. SRM mode was used to detect the target ions (m/z) of all derivatized metabolites and calibration standards. Metabolite concentrations were determined using calibration curves with internal standards.

Calibration standards and stable isotope-labeled (D, 13C) internal standards were purchased from the following sources:

- **Sigma-Aldrich (St. Louis, MO, USA):** γ-aminobutyric acid (GABA), L-alanine, L-leucine, L-leucine-5,5,5-D3, L-isoleucine, glycine.
- **Cambridge Isotopes (Tewksbury, MA, USA):** U-[13C]-GABA.

All standard solutions were prepared using ultra-pure water (Milli-Q). Derivatization reagents, including N-methyl-N-(trimethylsilyl)trifluoroacetamide with 1% trimethylchlorosilane (MSTFA + 1% TMCS), methoxyamine hydrochloride, and pyridine, were obtained from Sigma-Aldrich (St. Louis, MO, USA), while acetonitrile was purchased from J.T.Baker (Phillipsburg, NJ, USA). All reagents used had a purity of ≥98%.

Sample preparation for metabolite analysis was adapted from ^2^, with modifications. A 40 µL aliquot of each sample was mixed with 5 µL of internal standard (25 ng/µL: U-[13C]-GABA, L-leucine-5,5,5-D3) and 60 µL of acetonitrile using a vortex mixer for 10 min. The samples were then centrifuged at 14,000 × g for 4 min, and 40 μL of the supernatant was transferred to a new 1.5 mL GC-MS vial. The samples were dried using a Labconco Centrivap concentrator.

For derivatization, 50 μL of methoxyamine hydrochloride in pyridine (20 mg/mL) was added to each sample, followed by incubation at 37°C for 90 min with continuous mixing. Then, 70 μL of MSTFA was added, and the mixture was incubated at 37°C for 30 min. After an additional 2-hour equilibration at room temperature, 40 μL of acetonitrile was added, and the samples were mixed before GC-MS analysis.

1.	Pinu F., Villas-boas SG. Rapid Quantification of Major Volatile Metabolites in Fermented Food and Beverages Using Gas Chromatography-Mass Spectrometry. *Metabolites* 2017;**7**(3):37. Doi: 10.3390/metabo7030037.

2.	Sun S., Wang H., Xie J., Su Y. Simultaneous determination of rhamnose, xylitol, arabitol, fructose, glucose, inositol, sucrose, maltose in jujube (Zizyphus jujube Mill.) extract: comparison of HPLC–ELSD, LC–ESI–MS/MS and GC–MS. *Chemistry Central Journal* 2016;**10**(1):25. Doi: 10.1186/s13065-016-0171-2.
