## Supplementary material for "Dynamics and fermentation patterns of stool microbiota on simple carbon sources in *in vitro* batch cultures reveal dysbiosis in Crohn’s disease": Table S1: Table_S1.docx

**Table S1.** **Clinical characteristics of the CD patients, n = 7**

| **Value**  **Parameter** | **Average** | **Median** | **Min** | **Max** | **SD** |
| --- | --- | --- | --- | --- | --- |
| A. CD patients, n = 7 | | | | | |
| **Height (cm)** | 172.13 | 172.00 | 163.00 | 180.00 | 6.62 |
| **Weight (kg)** | 70.04 | 68.50 | 56.30 | 85.00 | 11.08 |
| **BMI (kg/m^2^)** | 23.59 | 22.75 | 19.38 | 30.12 | 3.30 |
| **HB (g/dl)** | 13.68 | 13.75 | 12.20 | 14.70 | 0.97 |
| **CRP (mg/l)** | 13.93 | 6.60 | 4.00 | 46.90 | 15.30 |
| **AST (U/L)** | 16.25 | 17.00 | 10.00 | 22.00 | 4.74 |
| **ALT (U/L)** | 16.00 | 14.00 | 8.00 | 31.00 | 7.60 |
| **GFR (mL/min/1.73 m^2^)** | 109.88 | 113.50 | 89.00 | 121.00 | 11.33 |
| **Creatinine (mg/dl)** | 0.76 | 0.81 | 0.50 | 0.87 | 0.12 |

SD - standard deviation, Fe – Iron, CRP – C Reactive Protein, AST – aspartate aminotransferase, ALT - alanine aminotransferase, HB – hemoglobin, GFR – glomerular filtration rate, BMI – Body Mass Index

**The following inclusion criteria for CD patients** were employed: (i) clinically-, endoscopically- and histopathologically-confirmed Crohn’s disease at least three months before the study; (ii) clinical symptoms confirmed by the attending physician; (iii) informed and voluntary consent to participate in the study; (iv) no antibiotic therapy and probiotic intake in the three months prior to the study; (v) no contraindications to blood donation.

**The following inclusion criteria for healthy individuals** were employed: (i) no medical history of disease symptoms; (ii) informed and voluntary consent to participate in the study; (iii) no antibiotic therapy and probiotic intake in the three months prior to the study; (iv) no contraindications to blood donation.
