## Supplementary material for "Dynamics and fermentation patterns of stool microbiota on simple carbon sources in *in vitro* batch cultures reveal dysbiosis in Crohn’s disease": Table S6: Table_S6.docx

| Table S6. Bacterial species showing statistically significant changes in the fecal microbiota of patients with Crohn's disease or in CD-stool-derived microbial communities. Table performed based on the data in Figure 4a. | |
| --- | --- |
| Species | **Short characteristics** |
| Species reduced in CD stool microbiotas only | |
| *Phocaeicola vulgatus*  (formerly *Bacteroides vulgatus*) | regarded as a relevant member of the core human microbiome that produces acetate, succinate, formate, propionate and other valuable metabolites such as 3-hydroxyphenylacetic acid, which helps maintain lipid homeostasis in the liver and prevents metabolic dysfunction-associated steatotic liver disease ^1,2^ |
| Species reduced both in CD stool microbiotas and CD-derived microbial communities grown on glucose* and/or acetate+lactate^#^ | |
| *Coprococcus catus* *^#^ | mainly a propionate- but also a butyrate-producer in the human microbiome, capable of lactate utilization  in the gut^3,4^ |
| *Ruminococcus torques* *^#^ | butyrate producer with decreased abundance in the microbiomes of CD patients,  especially those who are C-reactive protein (CRP) positive ^5^ |
| *Gemmiger formicilis* *^#^ | butyrate producer, reduced in both CD and colorectal cancer patients ^6^ |
| *Oscillibacter* sp. *^#^ | reduced in IBD patients ^7^ |
| *Anaerotignum faecicola* * | a bacterium isolated from human stool that produces acetate, propionate, butyrate and isobutyrate as fermentation end products ^8^, with no recognized role in IBD |
| *Alistipes putredinis* * | its role in the gut microbiome and IBD has yet to be confirmed by experimental studies ^9^. A human *A. putredinis* strain exhibited anti-inflammatory effects in a mouse model of colitis induced by sodium dextran sulfate  and oxazolone ^10^ |
| *Lactococcus lactis* ^#^ | a well-characterized bacterium with anti-inflammatory properties, commonly used as a probiotic ^11^ |
| Species reduced in CD-derived microbial communities grown on glucose and acetate+lactate | |
| *Fusicatenibacter saccharivorans* | producer of C2-C6 organic acids, including butyrate, in the human gut, with a decreased contribution to the gut microbiome of patients with colorectal cancer ^12^ and UC ^13^ |
| *Eubacterium rectale* (currently *Agathobacter rectalis*) | a well-recognized butyrate producer in the gut, reduced in CD and UC patients ^14–16^ |
| *Dorea longicatena*  *D. formicigenerans* | non-butyrate producers that ferment carbohydrates mainly to formate, acetate and ethanol ^17^, helping to maintain a healthy gastrointestinal environment by the downregulation of inflammation in the gut mucosa ^18^ *D. longicatena* was found to contribute to CD remission ^19^, whereas *D. formicigenerans* protects against colorectal carcinogenesis ^20^ |
| *Blautia faecis* | representing the *Blautia* genus, found in stool samples collected from a healthy human and regarded as a functional genus with potential probiotic properties due to its production of acetate, succinate, lactate, ethanol plus other beneficial effects for the host ^21^ |
| Species reduced in CD-derived microbial communities grown on acetate+lactate | |
| *Faecalibacterium prausnitzii* | a key beneficial commensal butyrate producer, with a well-documented diminished contribution in CD patients ^22–25^ |
| *Faecalibacillus intestinalis* | isolated from faeces of healthy humans, with potential probiotic properties ^26,27^ |
| *Anaerostipes hadrus* | an acetate-utilizing butyrate producer ^28^ |
| Species reduced in CD-derived microbial communities grown on glucose | |
| *Bifidobacterium adolescentis* | primary starch degrader and producer of acetate and lactate, stimulates the growth of butyrogenic bacteria through metabolic cross-feeding; the health-beneficial properties of B. adolescentis include enhancement of intestinal barrier function, anti-inflammatory and immune-regulatory effects, as well as the production of neurotransmitters (GABA) and vitamins. A reduced abundance of B. adolescentis in the human gut microbiome has been associated with various forms of IBD ^29–31^ |
| *Roseburia faecis*  (currently *Agathobacter faecis*) | a commensal bacterium that produces C2-C6 organic acids, particularly butyrate, which affects colonic motility, maintains immunity and has anti-inflammatory properties ^32^ |
| Species enriched in CD stool microbiotas only | |
| *Bifidobacterium bifidum* | the increases may result from disruption of the microbiome; the reduction in butyrate producers in CD patients appears to disturb the balance between different physiological groups of bacteria, leading to greater representation of members of the genus Bifidobacterium, for example ^33^ |
| Species enriched in CD-derived microbial communities grown on glucose | |
| *Escherichia coli* | well-documented pathobiont contributing to IBD and cancer ^34–37^ |
| *Enterococcus faecalis* | regarded as a probiotic strain with anti-inflammatory and protective properties; however, there are reports suggesting that it can cause injury to intestinal epithelial cells, induce gut inflammation, and promote cancer ^38,39^ |
| Species enriched in CD-derived microbial communities grown on acetate and lactate | |
| *Peptoniphilus harei* | regarded as a commensal in the human microbiome and also an opportunistic pathogen ^40,41^ |

References

1.	Jin S., Chen P., Yang J., Li D., Liu X., Zhang Y., et al. *Phocaeicola vulgatus* alleviates diet-induced metabolic dysfunction-associated steatotic liver disease progression by downregulating histone acetylation level via 3-HPAA. *Gut Microbes* 2024;**16**(1):2309683. Doi: 10.1080/19490976.2024.2309683.

2.	Clausen U., Vital S-T., Lambertus P., Gehler M., Scheve S., Wöhlbrand L., et al. Catabolic Network of the Fermentative Gut Bacterium *Phocaeicola vulgatus* (Phylum Bacteroidota) from a Physiologic-Proteomic Perspective. *Microb Physiol* 2024;**34**(1):88–107. Doi: 10.1159/000536327.

3.	Xu L., Yu Q., Ma L., Su T., Zhang D., Yao D., et al. In vitro simulated fecal fermentation of mixed grains on short-chain fatty acid generation and its metabolized mechanism. *Food Research International* 2023;**170**:112949. Doi: 10.1016/j.foodres.2023.112949.

4.	Sheridan PO., Louis P., Tsompanidou E., Shaw S., Harmsen HJ., Duncan SH., et al. Distribution, organization and expression of genes concerned with anaerobic lactate utilization in human intestinal bacteria. *Microbial Genomics* 2022;**8**(1). Doi: 10.1099/mgen.0.000739.

5.	Takahashi K., Nishida A., Fujimoto T., Fujii M., Shioya M., Imaeda H., et al. Reduced Abundance of Butyrate-Producing Bacteria Species in the Fecal Microbial Community in Crohn’s Disease. *Digestion* 2016;**93**(1):59–65. Doi: 10.1159/000441768.

6.	Ning L., Zhou Y-L., Sun H., Zhang Y., Shen C., Wang Z., et al. Microbiome and metabolome features in inflammatory bowel disease via multi-omics integration analyses across cohorts. *Nat Commun* 2023;**14**(1):7135. Doi: 10.1038/s41467-023-42788-0.

7.	Pisani A., Rausch P., Bang C., Ellul S., Tabone T., Marantidis Cordina C., et al. Dysbiosis in the Gut Microbiota in Patients with Inflammatory Bowel Disease during Remission. *Microbiol Spectr* 2022;**10**(3):e00616-22. Doi: 10.1128/spectrum.00616-22.

8.	Choi S-H., Kim J-S., Park J-E., Lee KC., Eom MK., Oh BS., et al. Anaerotignum faecicola sp. nov., isolated from human faeces. *J Microbiol* 2019;**57**(12):1073–8. Doi: 10.1007/s12275-019-9268-3.

9.	Parker BJ., Wearsch PA., Veloo ACM., Rodriguez-Palacios A. The Genus Alistipes: Gut Bacteria With Emerging Implications to Inflammation, Cancer, and Mental Health. *Front Immunol* 2020;**11**:906. Doi: 10.3389/fimmu.2020.00906.

10.	Ishikawa D., Zhang X., Nomura K., Shibuya T., Hojo M., Yamashita M., et al. Anti-inflammatory Effects of Bacteroidota Strains Derived From Outstanding Donors of Fecal Microbiota Transplantation for the Treatment of Ulcerative Colitis. *Inflammatory Bowel Diseases* 2024:izae080. Doi: 10.1093/ibd/izae080.

11.	Javid H., Oryani MA., Akbari S., Amiriani T., Ravanbakhsh S., Rezagholinejad N., et al. *L. Plantarum* and *L. Lactis* as a Promising Agent in Treatment of Inflammatory Bowel Disease and Colorectal Cancer. *Future Microbiol* 2023;**18**(16):1197–209. Doi: 10.2217/fmb-2023-0076.

12.	He T., Cheng X., Xing C. The gut microbial diversity of colon cancer patients and the clinical significance. *Bioengineered* 2021;**12**(1):7046–60. Doi: 10.1080/21655979.2021.1972077.

13.	Takeshita K., Mizuno S., Mikami Y., Sujino T., Saigusa K., Matsuoka K., et al. A Single Species of Clostridium Subcluster XIVa Decreased in Ulcerative Colitis Patients: *Inflammatory Bowel Diseases* 2016;**22**(12):2802–10. Doi: 10.1097/MIB.0000000000000972.

14.	Mukherjee A., Lordan C., Ross RP., Cotter PD. Gut microbes from the phylogenetically diverse genus *Eubacterium* and their various contributions to gut health. *Gut Microbes* 2020;**12**(1):1802866. Doi: 10.1080/19490976.2020.1802866.

15.	Rosero JA., Killer J., Sechovcová H., Mrázek J., Benada O., Fliegerová K., et al. Reclassification of Eubacterium rectale (Hauduroy et al. 1937) Prévot 1938 in a new genus Agathobacter gen. nov. as Agathobacter rectalis comb. nov., and description of Agathobacter ruminis sp. nov., isolated from the rumen contents of sheep and cows. *International Journal of Systematic and Evolutionary Microbiology* 2016;**66**(2):768–73. Doi: 10.1099/ijsem.0.000788.

16.	Pittayanon R., Lau JT., Leontiadis GI., Tse F., Yuan Y., Surette M., et al. Differences in Gut Microbiota in Patients With vs Without Inflammatory Bowel Diseases: A Systematic Review. *Gastroenterology* 2020;**158**(4):930-946.e1. Doi: 10.1053/j.gastro.2019.11.294.

17.	Taras D., Simmering R., Collins MD., Lawson PA., Blaut M. Reclassification of Eubacterium formicigenerans Holdeman and Moore 1974 as Dorea formicigenerans gen. nov., comb. nov., and description of Dorea longicatena sp. nov., isolated from human faeces. *International Journal of Systematic and Evolutionary Microbiology* 2002;**52**(2):423–8. Doi: 10.1099/00207713-52-2-423.

18.	Nishida K., Sawada D., Kawai T., Kuwano Y., Fujiwara S., Rokutan K. Para‐psychobiotic *Lactobacillus gasseri* CP 2305 ameliorates stress‐related symptoms and sleep quality. *J Appl Microbiol* 2017;**123**(6):1561–70. Doi: 10.1111/jam.13594.

19.	Mondot S., Lepage P., Seksik P., Allez M., Tréton X., Bouhnik Y., et al. Structural robustness of the gut mucosal microbiota is associated with Crohn’s disease remission after surgery. *Gut* 2016;**65**(6):954–62. Doi: 10.1136/gutjnl-2015-309184.

20.	Zhang X., Yu D., Wu D., Gao X., Shao F., Zhao M., et al. Tissue-resident Lachnospiraceae family bacteria protect against colorectal carcinogenesis by promoting tumor immune surveillance. *Cell Host & Microbe* 2023;**31**(3):418-432.e8. Doi: 10.1016/j.chom.2023.01.013.

21.	Liu X., Mao B., Gu J., Wu J., Cui S., Wang G., et al. *Blautia* —a new functional genus with potential probiotic properties? *Gut Microbes* 2021;**13**(1):1875796. Doi: 10.1080/19490976.2021.1875796.

22.	Gonzalez CG., Mills RH., Zhu Q., Sauceda C., Knight R., Dulai PS., et al. Location-specific signatures of Crohn’s disease at a multi-omics scale. *Microbiome* 2022;**10**(1):133. Doi: 10.1186/s40168-022-01331-x.

23.	Quévrain E., Maubert MA., Michon C., Chain F., Marquant R., Tailhades J., et al. Identification of an anti-inflammatory protein from *Faecalibacterium prausnitzii* , a commensal bacterium deficient in Crohn’s disease. *Gut* 2016;**65**(3):415–25. Doi: 10.1136/gutjnl-2014-307649.

24.	Leylabadlo HE., Ghotaslou R., Feizabadi MM., Farajnia S., Moaddab SY., Ganbarov K., et al. The critical role of Faecalibacterium prausnitzii in human health: An overview. *Microbial Pathogenesis* 2020;**149**:104344. Doi: 10.1016/j.micpath.2020.104344.

25.	Lenoir M., Martín R., Torres-Maravilla E., Chadi S., González-Dávila P., Sokol H., et al. Butyrate mediates anti-inflammatory effects of *Faecalibacterium prausnitzii* in intestinal epithelial cells through *Dact3*. *Gut Microbes* 2020;**12**(1):1826748. Doi: 10.1080/19490976.2020.1826748.

26.	Zhang Z., Mocanu V., Deehan EC., Hotte N., Zhu Y., Wei S., et al. Recipient microbiome-related features predicting metabolic improvement following fecal microbiota transplantation in adults with severe obesity and metabolic syndrome: a secondary analysis of a phase 2 clinical trial. *Gut Microbes* 2024;**16**(1):2345134. Doi: 10.1080/19490976.2024.2345134.

27.	Seo B., Jeon K., Baek I., Lee YM., Baek K., Ko G. Faecalibacillus intestinalis gen. nov., sp. nov. and Faecalibacillus faecis sp. nov., isolated from human faeces. *International Journal of Systematic and Evolutionary Microbiology* 2019;**69**(7):2120–8. Doi: 10.1099/ijsem.0.003443.

28.	Endo A., Tanno H., Kadowaki R., Fujii T., Tochio T. Extracellular fructooligosaccharide degradation in Anaerostipes hadrus for co-metabolism with non-fructooligosaccharide utilizers. *Biochemical and Biophysical Research Communications* 2022;**613**:81–6. Doi: 10.1016/j.bbrc.2022.04.134.

29.	Leser T., Baker A. Bifidobacterium adolescentis – a beneficial microbe. *Benef Microbes* 2023;**14**(6):525–51. Doi: 10.1163/18762891-20230030.

30.	Kowalska-Duplaga K., Gosiewski T., Kapusta P., Sroka-Oleksiak A., Wędrychowicz A., Pieczarkowski S., et al. Differences in the intestinal microbiome of healthy children and patients with newly diagnosed Crohn’s disease. *Sci Rep* 2019;**9**(1):18880. Doi: 10.1038/s41598-019-55290-9.

31.	Duranti S., Ruiz L., Lugli GA., Tames H., Milani C., Mancabelli L., et al. Bifidobacterium adolescentis as a key member of the human gut microbiota in the production of GABA. *Sci Rep* 2020;**10**(1):14112. Doi: 10.1038/s41598-020-70986-z.

32.	Tamanai-Shacoori Z., Smida I., Bousarghin L., Loreal O., Meuric V., Fong SB., et al. *Roseburia* Spp.: A Marker of Health? *Future Microbiol* 2017;**12**(2):157–70. Doi: 10.2217/fmb-2016-0130.

33.	Wang W., Chen L., Zhou R., Wang X., Song L., Huang S., et al. Increased Proportions of Bifidobacterium and the Lactobacillus Group and Loss of Butyrate-Producing Bacteria in Inflammatory Bowel Disease. *J Clin Microbiol* 2014;**52**(2):398–406. Doi: 10.1128/JCM.01500-13.

34.	Sugihara K., Kamada N. Metabolic network of the gut microbiota in inflammatory bowel disease. *Inflamm Regener* 2024;**44**(1):11. Doi: 10.1186/s41232-024-00321-w.

35.	Mirsepasi-Lauridsen HC., Vallance BA., Krogfelt KA., Petersen AM. *Escherichia coli* Pathobionts Associated with Inflammatory Bowel Disease. *Clin Microbiol Rev* 2019;**32**(2):e00060-18. Doi: 10.1128/CMR.00060-18.

36.	Pobeguts OV., Ladygina VG., Evsyutina DV., Eremeev AV., Zubov AI., Matyushkina DS., et al. Propionate Induces Virulent Properties of Crohn’s Disease-Associated Escherichia coli. *Front Microbiol* 2020;**11**:1460. Doi: 10.3389/fmicb.2020.01460.

37.	Ormsby MJ., Logan M., Johnson SA., McIntosh A., Fallata G., Papadopoulou R., et al. Inflammation associated ethanolamine facilitates infection by Crohn’s disease-linked adherent-invasive Escherichia coli. *EBioMedicine* 2019;**43**:325–32. Doi: 10.1016/j.ebiom.2019.03.071.

38.	Quaglio AEV., Grillo TG., Oliveira ECSD., Stasi LCD., Sassaki LY. Gut microbiota, inflammatory bowel disease and colorectal cancer. *WJG* 2022;**28**(30):4053–60. Doi: 10.3748/wjg.v28.i30.4053.

39.	De Almeida CV., Taddei A., Amedei A. The controversial role of *Enterococcus faecalis* in colorectal cancer. *Therap Adv Gastroenterol* 2018;**11**:1756284818783606. Doi: 10.1177/1756284818783606.

40.	Wan X., Wang S., Wang M., Liu J., Zhang Y. Identification of Peptoniphilus harei From Blood Cultures in an Infected Aortic Aneurysm Patient: Case Report and Review Published Literature. *Front Cell Infect Microbiol* 2021;**11**:755225. Doi: 10.3389/fcimb.2021.755225.

41.	Diop K., Diop A., Michelle C., Richez M., Rathored J., Bretelle F., et al. Description of three new *Peptoniphilus* species cultured in the vaginal fluid of a woman diagnosed with bacterial vaginosis: *Peptoniphilus pacaensis* sp. nov., *Peptoniphilus raoultii* sp. nov., and *Peptoniphilus vaginalis* sp. nov. *MicrobiologyOpen* 2019;**8**(3):e00661. Doi: 10.1002/mbo3.661.
